# ND-13, a DJ-1 derived peptide, as a novel pharmacological approach in the prevention of NLRP3 inflammasome activity in Diabetic Nephropathy

**DOI:** 10.1101/2025.01.21.633908

**Authors:** María José Caballero-Herrero, Celia Arias-Sánchez, Ignacio Quevedo, Patricia S. Latham, Carmen De Miguel, Yihan Zhong, Gaoxi Xu, Estela Guillen, Adrián Núñez-Sancho, Diego Angosto, Cristina Molina-López, Laura Hurtado-Navarro, Julieta Schachter, Luis Pardo Marín, José Joaquín Cerón, Antonio M Hernandez-Martinez, Florentina Rosique López, Leonor Andúgar Rocamora, Juan B Cabezuelo, Pablo Pelegrin, Santiago Cuevas

## Abstract

Diabetic nephropathy is the most important cause of renal failure worldwide and is characterized by sustained inflammation regulated in part by NLRP3 Inflammasome. Attenuation of inflammation is a major priority to prevent renal damage. Our previous publications show that DJ-1 has antioxidant and anti-inflammatory properties in the kidney. ND-13 is a short peptide consisting of 13 amino acids of the DJ-1-protein, which could increase DJ-1 pathway activation. The aim of these studies was to determine the role of NLRP3 in the pathogenesis of diabetic nephropathy and to study the possible renal protective effects of the DJ-1 pathway in diabetic mice and on inflammasome regulation.

Bone marrow derived macrophages were treated with ND-13 and cultivated in high and low glucose. Diabetes was induced in C57Bl/6 mice via injection of streptozotocin and treated with ND-13 and MCC950, an inhibitor of the NLRP3 inflammasome. Peripheral mononuclear cells were isolated from human with diabetes, diabetic nephropathy and healthy donors and were plated, pretreated with ND-13 and stimulated with LPS+ATP.

IL-1β concentration in the medium of bone marrow derived macrophages increased by NLRP3 inflammasome stimulation by LPS+ATP, and decreased in macrophages pre-treated with ND-13, however, in the presence of LPS+Nigericin no effect was found. Peritoneal macrophages from diabetic mice were obtained and plated. Streptozotocin-induced diabetic C57BL/6 mice have increased the peritoneal cells IL-1β production compared with control mice, suggesting that inflammasome may be activated in macrophages during diabetes and that ND-13 treatment normalized its activity. ND-13 and MCC950 decreased histologic evidence of tubular injury in STZ-induced diabetes in mice, additionally, significantly increased the mRNA expression of *Col-I, Col-II, Tgf-β, Il-6, Tnf-α* and *P2x7* in the renal cortex, which was partially prevented by ND-13 and MCC950 pre-treatment. *P2x7* mRNA expression was also increased in peritoneal macrophages obtained from diabetic mice, and its expression was attenuated by ND-13 pre-treatment. Peripheral mononuclear cells isolated from patient’s blood were plated and stimulated with LPS+ATP. Patients with diabetic nephropathy presented a significant increase of IL-1β release compared to diabetic individuals and ND-13 could has a role in the prevention of inflammasome activation in healthy patients.

Our results demonstrate that activation of the DJ-1 pathway is a promising approach to prevent renal inflammation and fibrosis during diabetes, by ameliorating inflammasome activation in peripheral immune cells. Thus, ND-13 could be a promising new therapeutic approach to attenuating inflammation and renal damage in diabetic nephropathy.

## 1. Introduction

Diabetic nephropathy (DN) is the most common cause of renal failure worldwide and diabetes and hypertension are the main causes of chronic kidney disease (CKD) [1]. The current diagnosis of CKD is estimated by glomerular filtration rate <60 mL/min/1.73 m^2^ and an elevated urinary albumin-excretion rate is a known early predictor of future cardiovascular events [2]. Renal oxidative stress and inflammation are two of the most important factors involved in the pathogenesis of DN and other cardiovascular complications [3]. Currently, the development of renal disease in diabetes patients cannot be prevented with the current pharmacological therapies, therefore, new approaches to therapy and targets are urgently required [4].

It is known that kidney disease is characterized by progressive destruction of the renal parenchyma by sustained inflammation [5, 6], and attenuating inflammation and fibrosis in these patients is the major priority to prevent renal damage. Inflammation and the consequent oxidative stress, and *vice versa*, are considered major factors triggering fibrosis and key components in the development and progression of renal failure [5]. Renal inflammation plays a central role in the initiation and progression of fibrosis in kidney disease; therefore, the attenuation of the inflammatory response may be a critical step for the restoration of the proper balance between pro- and anti-fibrotic signaling pathways [7] in the kidney and other organs.

Inflammasomes are central players in the inflammatory response and represent one of the initial steps for the induction of renal inflammation (reviewed in [8]). Inflammasomes are considered intracellular sensors for pathogens and tissue damage [9]. The sensor protein gives the name to the inflammasome and shares similar structural domains, belonging to the nucleotide-binding domain and leucine-rich-containing family of receptors (NLRs). NLRP3 is the most studied inflammasome [10, 11] because it is considered as the main trigger of pathogen-free inflammation (**Figure 1**) and has been implicated in the pathophysiology of different autoinflammatory syndromes [12] as well as numerous diseases associated with inflammatory, metabolic, and degenerative processes, including renal diseases [8, 13]. Given the central role of NLRP3 activation in the pathogenesis of diabetes[14], targeting the NLRP3 inflammasome presents a promising therapeutic strategy. Therapies that neutralize IL-1β or block its receptor have shown promise in reducing inflammation and improving glycemic control in diabetic patients. A recent meta-analysis shown that anti-inflammatory compounds targeting interleukin-1β (IL-1β), interleukin-1β receptor (IL-1βR), tumor necrosis factor-α (TNF-α), and nuclear factor-κB (NF-κB) significant reduce of the levels of fasting plasma glucose (FPG), glycosylated haemoglobin (HbA1c), and c-reactive protein (CRP) in patients with Type 2 diabetes compared with control, and combination therapies targeting IL-1β and TNF-α improved beneficial effects than targeting IL-1β or TNF-α alone [15], therefore, small molecules or biological agents that directly inhibit NLRP3 activation or its downstream signaling pathways may have a good approach to prevent the morbidity in diabetes.

**Figure 1.**
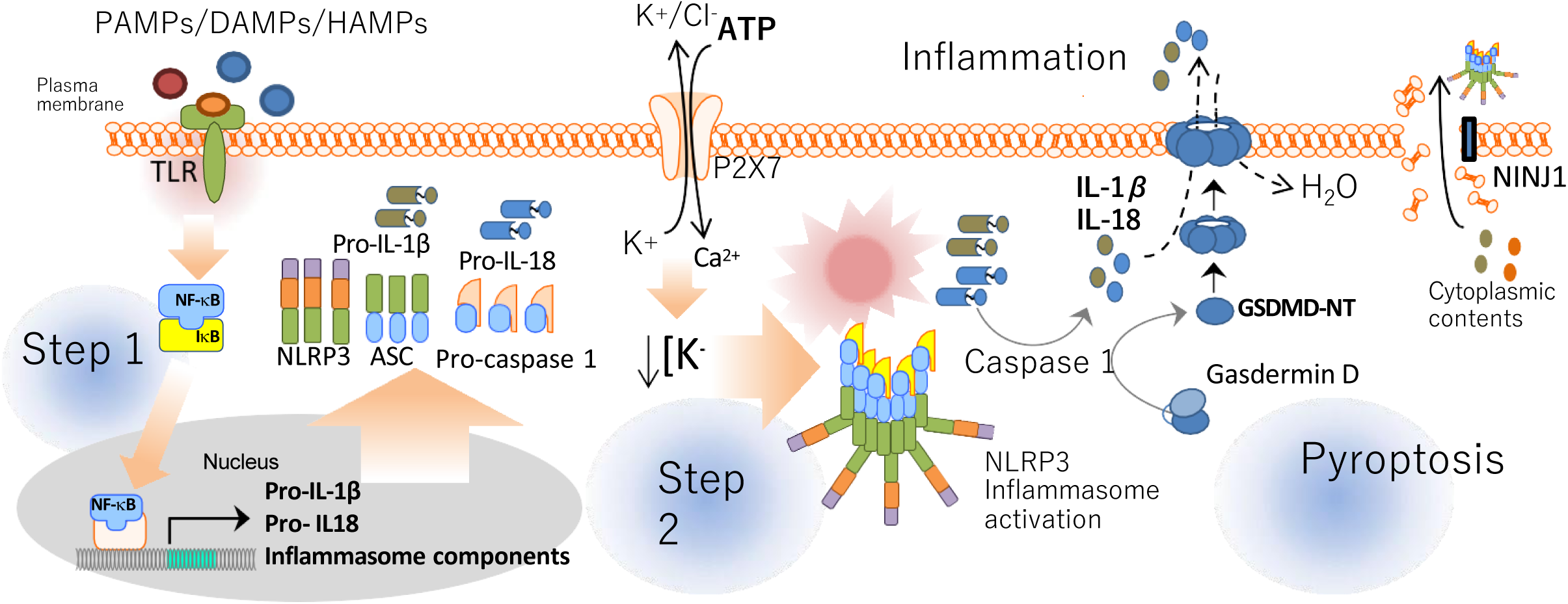
Activation of NLRP3 inflammasome. NLRP3 inflammasome needs two consecutive steps to be active: a priming stage and an activation stage. During the priming, TLR receptors can be stimulated by DAMPs, HAMPs and PAMPs which allow the translocation of nuclear factor kappa B (NF-κB) to the cell nucleus. The activation of NF-κB pathway increase the transcription of the inflammasome components, and the immature form of some pro-inflammatory cytokines. Next, a second signal such as P2X7 activation by extracellular ATP, lysosomal destabilization and/or production of ROS is required to induces the oligomerization of NLRP3 inflammasome complex which activate caspase 1. Caspase 1 can turn pro-IL-1β and pro-IL-18 into mature cytokines and cleaves the N-terminus of Gasdermin D which can oligomerizes in membranes to form pores. Liberation of pro-inflammatory cytokines such as IL-1β and IL-18, NLRP3 inflammasome complex and other cytoplasmic contents induces inflammation and a cell death type called pyroptosis. (Figure modified from [71]).

DJ-1 (also known as Park 7), which was initially identified as an autosomal recessive gene associated with Parkinson’s disease, is expressed in brain, heart, kidney, liver, pancreas, and skeletal muscle in rodents and humans [16]. DJ-1 is a multifunctional oxidative stress response protein that functions as a redox-sensitive chaperone with intrinsic antioxidant properties, especially in the mitochondria, and regulates the expression of several antioxidant genes [17–19]. We have reported that the renal *DJ-1* has renal antioxidant and anti-inflammatory properties [20, 21], mice with *DJ-1* selectively silenced in the kidney and mice with germline deletion of *DJ-1* (*DJ-1^-/-^)* have high blood pressure (BP), and decreased expression and activity of Nrf2 [20], suggesting that DJ-1 can inhibit renal reactive oxygen species (ROS) production, at least in part, via the activation of Nrf2-antioxidant genes. Moreover, it has been reported that DJ-1 could be an interesting target in the pathogenesis of diabetic nephropathy in rats [22].

Dr. Offen’s laboratory has developed a peptide from the amino acid sequence of DJ-1. The active areas of DJ-1 were studied, and the most conserved area was selected; a 13 amino acid chain from DJ-1. To achieve cell permeability, the 13 amino acid chain was linked to a 7-amino acid cell-penetrating peptide. The resulting 20 amino acid compound was named: ND-13. It has been repeatedly demonstrated that neuronal cultures treated with ND-13 are protected from the effects of exposure to Parkinson’s disease (PD)-relevant neurotoxins and in other pathologies [23–25]. ND-13 reduces apoptosis and inactivates the pro-apoptotic caspase-3 in neuronal cell lines exposed to neurotoxic insults. The cells treated with ND-13 activated the Nrf2 pathway, resulting in the increased expression of Nrf2-induced antioxidant genes, such as Heme-Oxygenase-I (*Hmox1*), *Nqo1, Gclc*, and *Gclm*, similar to our findings in the kidney [20]. Our previous publication demonstrates that ND-13 prevent inflammation and have protective effects on a renal damage model induced by effect in unilateral ureter obstruction [26].

Despite the growing evidence of the involvement of NRLP3 inflammasome in diabetic kidney disease and the kidney protective effects of DJ-1 pathway activation, it is still unclear if there is a connection that involves DJ-1 pathway activation and NLRP3 inflammasome activation that could ameliorate kidney disease during diabetes. So, the main goal of this study is to determine the role of NLRP3 inflammasome activation in patients with DN and study the possible effects of ND-13 on inflammasome activation and in prevention of renal damage in diabetes.

## 2. Results

### 2.1 ND-13 treatment reduces Il-1β release in BMDM but not TNF-α release

ND-13 has been described as a potential therapeutic approach to prevent renal disease thanks to his anti-inflammatory and anti-fibrotic properties found in an UUO model (Unilateral ureter obstruction model) [26]. To better understand his anti-inflammatory properties, we evaluated his implication or not in the activation of NLRP3 inflammasome, an important link between inflammation and renal damage [27, 28]. We used LPS as signal priming and two different activators of NLRP3 inflammasome, Adenosine triphosphate (ATP) and Nigericin. Activation of NLRP3 inflammasome was measured by release of Il-1β in the culture medium and we found a significant decrease in BMDM activated with LPS+ATP and treated with ND-13 compared with BMDM activated with same stimulus and no treatment (348.3±8.495 vs 197.0±45.19, n=3, t-test, *p< 0.05) (**Figure 2a**). No significant differences were found for BMDM treated or not with ND-13 and activated with LPS+Nigericin furthermore, TNF-α release was also not significant after LPS stimulus (**Figure 2a-b**). There was also no significant the cell death between the different stimuli or compare with resting results (**Figure 2c**). These data suggest that ND-13 could act by reducing IL-1β release in BMDM and therefore NLRP3 inflammasome activation but not in the activation of NF-κB pathway. Differences found between both NLRP3 activators (ATP and Nigericin) may be due to the fact that each compound induces NLRP3 inflammasome activation by different mechanisms.

**Figure 2.**
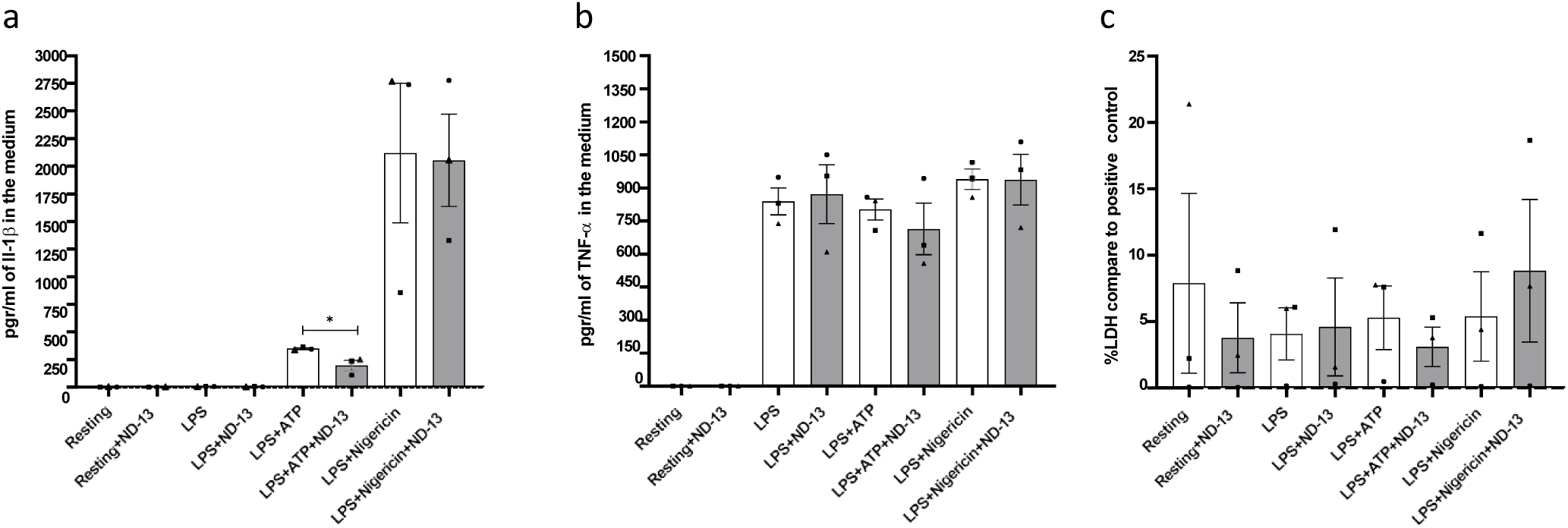
ND-13 prevents NLRP3 inflammasome activation in BMDM but not affect NF-κB pathway activation. **(a)** Release of Il-1β in the culture medium. **(b)** Release of TNF-α in the culture medium. **(c)** Percentage of cell death compared to positive control. Data is represented as mean ± SEM (n=3); t-test two-sided was used in a-c to compare between the group without ND-13 (white column) and its corresponding group with ND-13 (grey column); *\*p < 0.05*.

### 2.2 ND-13 treatment reduces NLRP3 inflammasome activation in diabetic mice

It has been described that hyperglycaemia can induce activation of NLRP3 inflammasome by TXNIP, a prooxidative and proinflammatory protein, with the consequent production of renal damage [29] so we wanted to evaluate ND-13 anti-inflammatory properties in diabetic nephropathy as a promising pharmacological approach for this renal disease. However, we could not see a difference in NLRP3 inflammasome activation comparing BMDM cultivated in high glucose with BMDM cultivated in low glucose, possibly due that multiple factors may involve in the high glucose-induced inflammation in human [30](**Supplementary figure 3a**). By contrast, we obtained a significant increase of TNF-α release in BMDM cultivated in low glucose compare with BMDM cultivated in high glucose (**Supplementary figure 3b**). It is known that hypoglycaemia primes the innate immune system and increases the inflammatory response to a subsequent inflammatory stimulus, which may explain these results, thus, this could make it difficult to test our hypothesis in vitro [31, 32]. Cell death was also not significant comparing resting, LPS+ATP or LPS+Nigericin between both condition (**Supplementary figure 3c**) suggesting that pyroptosis was not affected by glucose concentrations.

In order to pursue our objective of determining the anti-inflammatory effects of ND-13 in diabetic nephropathy, we created STZ-induced diabetic C57BL/6 mice as a model of diabetes and we treat some of them with ND-13 or MCC950, a specific NLRP3 inflammasome inhibitor [33]. Control group, that means no diabetic C57BL/6 mice, were treated with citrate buffer and showed higher weight and less blood-glucose levels than its littermates. STZ-induced diabetic C57BL/6 mice showed a significant weight decrease compared with control mice (32.65±0.6516 vs 20.02±1.157; 24.69±1.396; 24.81±1.086, n=7-10, one-way ANOVA multiple comparisons, ****p<0.0001, ***p<0.0005, *p<0.05), especially in mice without treatment of ND-13 or MCC950. However, we could not see significant differences in blood-glucose levels during all the treatment process between the different STZ-induced diabetic C57BL/6 mice groups, which suggest that treat diabetic mice with ND-13 or MCC950 do not affect diabetic induction but could ameliorate weight loss, suggesting that the animals were in better health condition with this treatments despite having similar levels of hyperglycaemia (**Figure 3a-b**). Furthermore, we found that STZ-induced diabetic C57BL/6 mice groups without additional treatment had a significant increase in the urinary albumin-creatinine ratio compared to the control mice, indicating that these mice have affected glomerular filtration (**Figure 3c**). Next, we collected and stimulated in vitro peritoneal immune cells from all mice to evaluate NLRP3 inflammasome activation in this mouse model. Stimulation with LPS+ATP showed a large and significant increase of IL-1β release in STZ-induced diabetic C57BL/6 mice compared with control group (2120±368.7 vs 4476 ±428.6; n=7-10, t-test, **p < 0.01), however this significant increase did not occur in diabetic mice treated with ND-13 (4476 ±428.6 vs 2240± 382.4; n=7-10, t-test, **p < 0.01) or the NLRP3 inflammasome inhibitor (4476 ±428.6 vs 2887± 423.6; n=8-10, t-test, **p < 0.01) (**Figure 3d**). Strengthening previous literature [34], our data showed that NLRP3 inflammasome may be active in diabetes.

**Figure 3.**
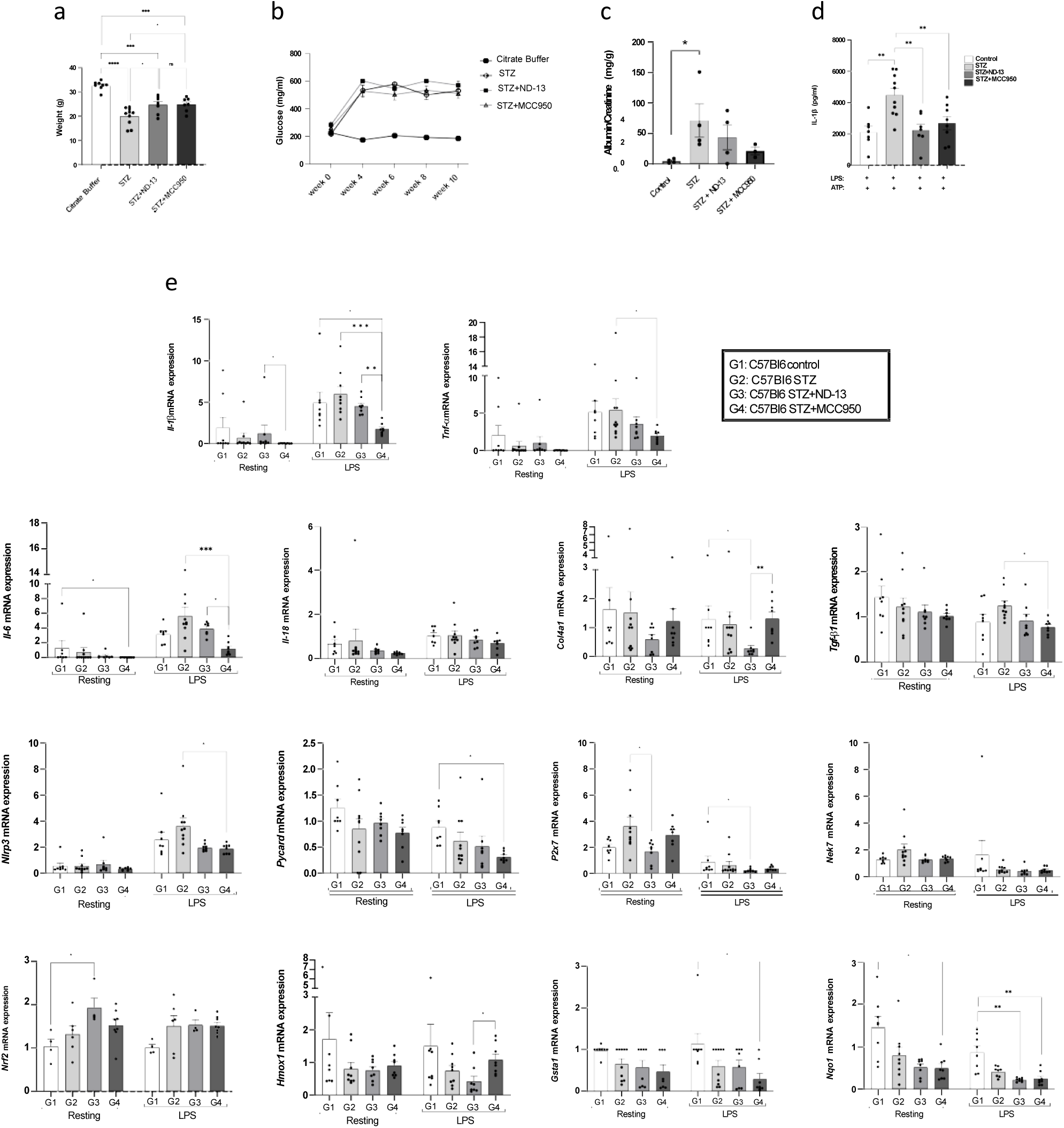
ND-13 reduces NLRP3 inflammasome activation in peritoneal macrophages cells from STZ-induced diabetic C57BL/6 mice. **(a)** Weight of mice at the end pf the treatment **(b)** Blood glucose levels of mice. **(c)** Urinary albumin-creatinine ratio in mice. **(d)** Effects of ND-13 and MCC950 on inflammasome activation in mouse peritoneal macrophages stimulated by LPS plus ATP. **(e)** mRNA expression of genes associated with fibrosis, inflammation, NLRP3 inflammasome activation and Nrf2 activation in mouse peritoneal macrophages cells. Data is represented as mean ± SEM; Kruskal-Wallis test or one-way ANOVA multiple comparisons was used in a and e, significance levels are indicated as follow *\*\*\*\*p<0.0001*, *\*\*\*p<0.001*, *\*\*p<0.01*, *\*p<0.05*, ns indicates no significant difference (*p>0.05*); t-test two-sided was used in d, p**<0,01. Control n=8, STZ n=10, STZ+ND-13 n=8, STZ+MCC950 n=8. one-way ANOVA was used in c to compare all groups to the control group, *p<0.05; Control n=4, STZ n=4, STZ+ND-13 n=4, STZ+MCC950 n=3.

We therefore evaluated mRNA expression searching for some proinflammatory, profibrotic or anti-inflammatory genes. mRNA expression from peritoneal cells without stimulus did not show significant differences between the different mice groups except for *Il-1β, Nrf2, Nq01, Il-6* and *P2x7*. *Nrf2* expression showed a significant increase in STZ-induced diabetic C57BL/6 mice treated with ND-13 compared with control group, to the contrary *Nqo1* expression tends to decrease in the three STZ-induced diabetic C57BL/6 mice groups comparing them with the expression of the control group, being significantly different in the case of mice treated with MCC950. Interestingly, we observed a significantly decrease in *P2x7* expression in diabetic mice treated with ND-13 compared with diabetic mice without treatment. The expression of *Nlrp3* was markedly elevated in all four groups of mice following stimulation with LPS, and the differences observed in the MCC950 group were notably greater in comparison to the other groups. The pro-inflammatory cytokines examined, *Il-1β, Il-18, Tnf-α,* and *Il-6*, demonstrated elevated expression levels following LPS stimulation, when compared to unstimulated cells within the four mouse groups. Profibrotic genes expression such as *Col4a1* significantly decreased in cells stimulated with LPS from mice treated with ND-13 compared with LPS stimulated cells from control group while *TGF-β* expression was significantly decreased in cells stimulated with LPS in mice treated with MCC950 comparing them with cells from the diabetic mice (Figures 3E).

### 2.3 STZ-induced diabetic C57BL/6 mice promote renal fibrosis and inflammation at the level of gene expression but not at tissue level

Activation of NLRP3 inflammasome is involved in the pathology of different kidney disease, including diabetic nephropathy [29, 35]. A characteristic of diabetic nephropathy is the loss of kidneys function causes by uncontrolled renal inflammation and fibrosis that in some cases ended in renal failure [36]. Therefore, to determine whether the peptide ND-13 could be a promising preventive treatment for diabetic patients, we evaluated renal damage in kidneys from control mice and STZ-induced diabetic C57BL/6 mice treated or not with ND-13 or MCC950. We did not find a significant difference in any of the four groups of mice in renal infiltration of macrophages or CD3+ cells even not in blue trichrome stain (**Figure 4**). The protein expression of desmin and synaptopodin were assayed by Western blot in whole renal tissue from control, STZ and STZ+ND-13 mice and showed no differences between the groups (**Supplementary figure 2a-b**) GSK3 and P-GSK3 also did not show significant differences between BMDM control and BMDM treated with ND-13 (**Supplementary figure 2c**) this indicates that incipient kidney inflammation and NRLP3 activation affect the kidneys, but kidney damage has not yet occurred, suggesting that these mechanisms play an important role in incipient kidney inflammation.

**Figure 4.**
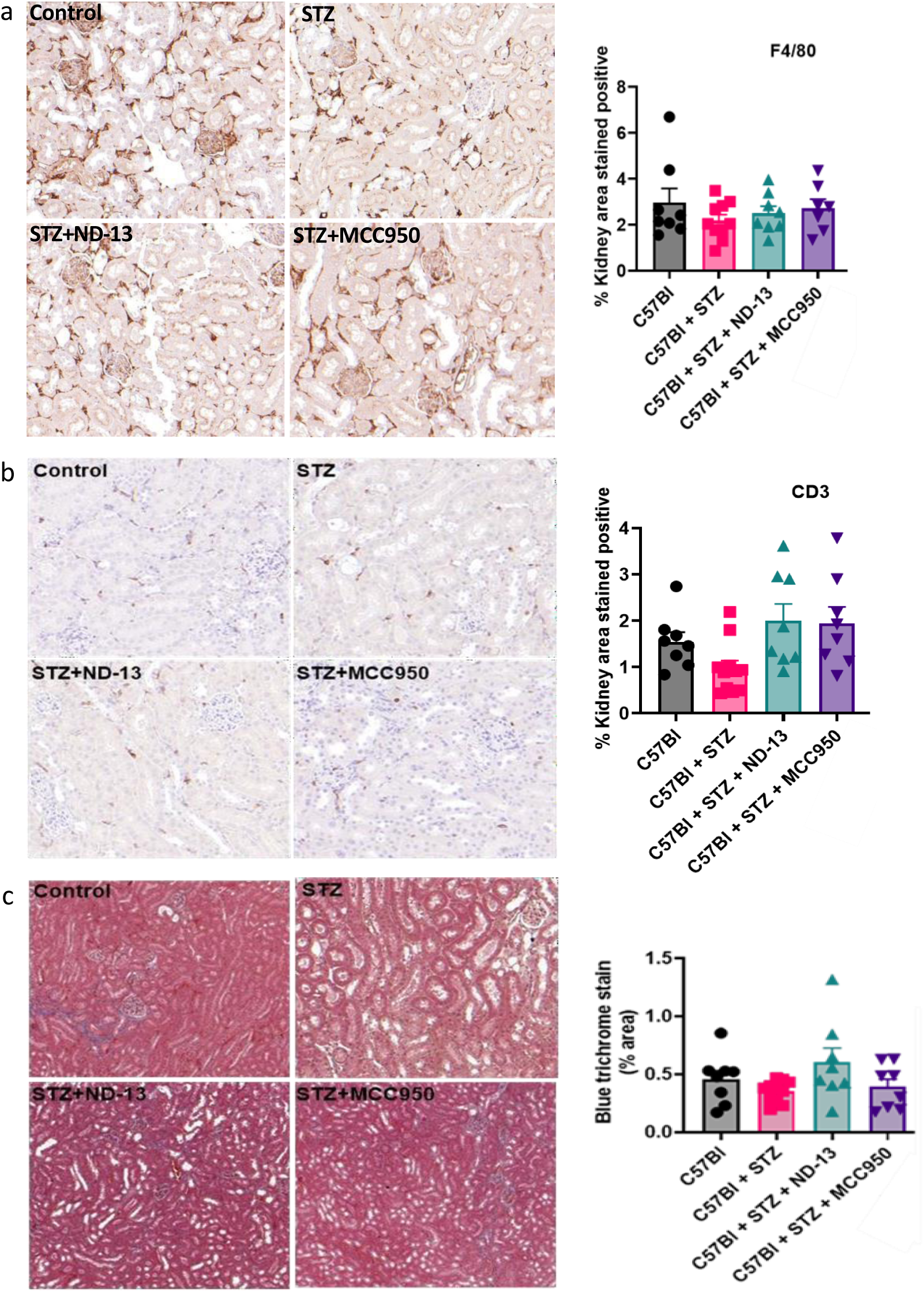
STZ-induced diabetic C57BL/6 mice do not promote renal infiltration of immune cells and renal fibrosis. **(a)** Infiltration of macrophages in the whole kidney. **(b)** Infiltration of T-cells in the whole kidney. **(c)** Fibrosis in the whole kidney. Quantification of infiltration and fibrosis was performed by taking 10 cortical images from each animal (n=7-8) at 400× magnification. The data are reported as mean ± SEM of positively stained area/glomerular area (%). The average percentage area that stained positively for F4/80 or CD3+ per experimental group. Control n=8, STZ n=10, STZ+ND-13 n=8, STZ+MCC950 n=8. No significant differences were found.

Whole renal tissue qPCR analyses showed a significant increased expression of *Tnf-α, Col1a1, P2x7, Pycard, Nq01 and Gta1* in STZ mice without treatment compared with control mice, significant increase that is lost in STZ mice treated with ND-13 or MCC950 (**Figure 5**). Taken together, these analyses indicates that our STZ-induced diabetic model cannot produce enough renal inflammation and fibrosis to determine whether ND-13 can reduce effectively renal inflammation causes by NLRP3 inflammasome and prevent renal damage in diabetic mice.

**Figure 5.**
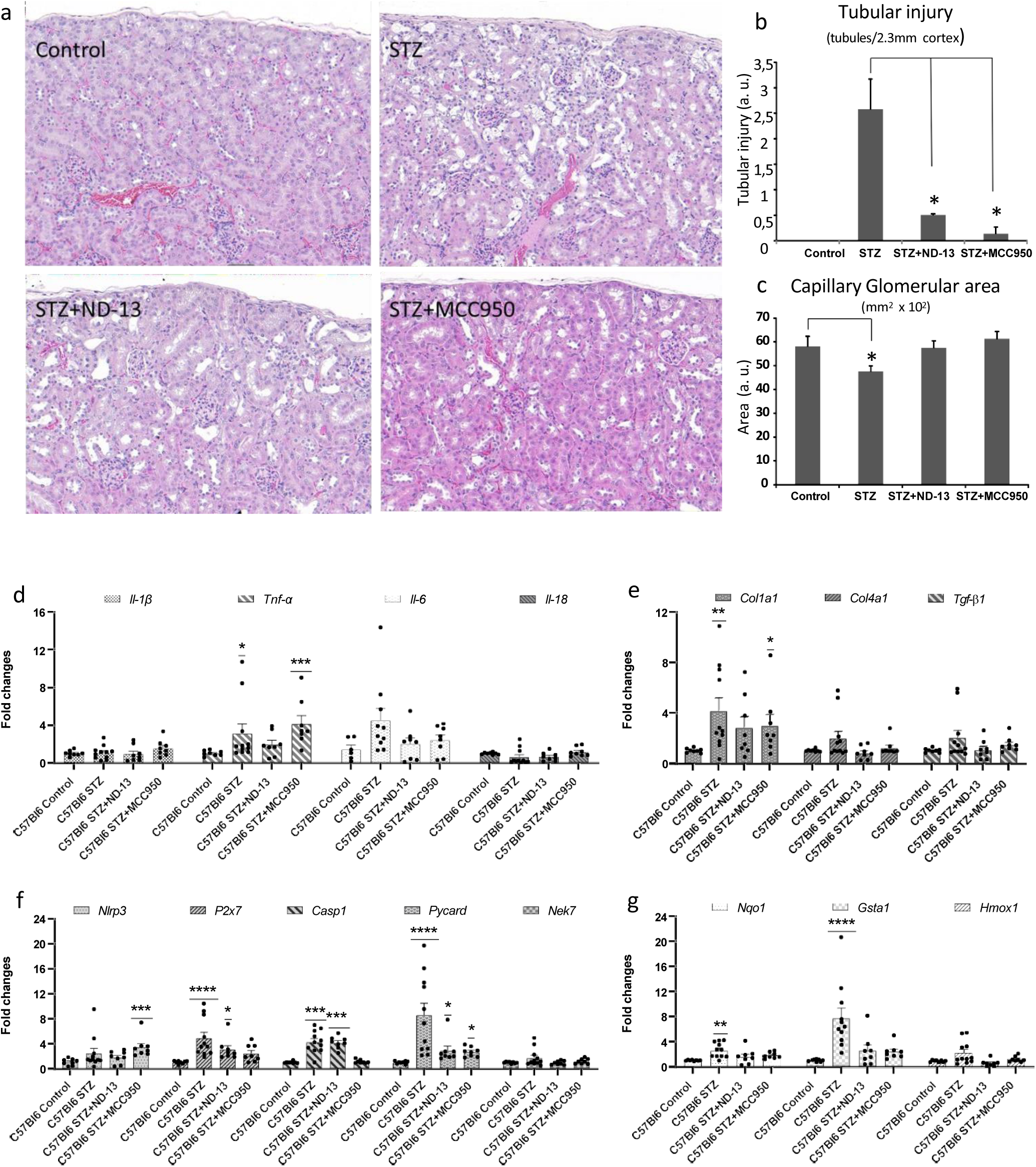
STZ-induced diabetic C57BL/6 mice promotes tubular damage, reduction of capillary glomerular area, renal fibrosis and inflammation at the level of gene expression. **(a)** Histologic sections of mouse renal cortex (100X) with H&E staining: Normal control mice, STZ-induced diabetes mice, STZ-induced diabetes mice with treatment of ND-13, STZ-induced diabetes mice with treatment of MCC950. **(b/c).** Tubular Injury and Glomerular Capillary area in normal and diabetic (STZ) mice with and without treatment. Data are the mean SD: Tubular injury of STZ vs control (*P = 0.0018*); STZ ND-13 vs STZ (*P = 0.0158*); STZ MCC950 vs STZ (*P = 0.0030*); Glomerular capillary area of STZ vs control (*P = 0.0245*); STZ ND-13 and STZ MCC950 vs control (*P = 0.8506* and *P = 0.5881*), respectively. **(d)** mRNA expression in the renal cortex of genes associated with inflammation**. (e)** mRNA expression in the renal cortex of genes associated with fibrosis. **(f)** mRNA expression in the renal cortex of genes associated with NLRP3 inflammasome activity. **(g)** mRNA expression in the renal cortex of antioxidant genes associated with Nrf2 activation. Data is represented as mean ± SEM in d-g; Dunn’s multiple comparisons test was used to compare each group with control group; significance levels are indicated as follow *\*\*\*\*p<0.0001*, *\*\*\*p<0.001*, *\*\*p<0.005*, *\*p<0.05*. Control n=8, STZ n=10, STZ+ND-13 n=8, STZ+MCC950 n=8. Arbitrary unit (a.u.).

On the other hand, P2x7 expression was decreased in STZ mice treated with ND-13 in whole tissue (**Figure 5f**) consistently with the trend observed in peritoneal cells on P2x7 expression (**Figure 3e**), these observations suggest that reducing P2x7 expression may be one of the protective mechanisms induced by ND-13.

### 2.4. STZ-induced diabetic C57BL/6 mice treated with ND-13 or MCC950 has less tubular injury and glomerular capillary area decreased than diabetic mice without treatment

The tubular and glomerular damage present in the kidneys obtained from the experimental mice were evaluated by Patricia S. Latham, MD, EdD in the Departments of Pathology and Internal Medicine of The George Washington University School of Medicine and Health Sciences, using hematoxylin and eosin (H&E) stained sections. Diabetic (STZ-treated) mice showed evident dilation of the renal pelvis and generalized, patchy dilation of tubules and Bowman’s space with occasional epithelial cell glycogen nuclei. A significant increase in tubular injury was seen in the cortex with focal attenuation of tubules and focal epithelial cell ballooning and dropout of the lining epithelial cells. Treatment of diabetic mice with ND-13 or MCC950 resulted in a significant decrease in evidence of tubular injury. Proteinaceous material was found in the tubules, but there were no dense deposits to suggest cast formation. There were occasional, small parenchymal clusters of mononuclear cells (< 10 cells) that were commonly seen in both diabetic and normal control mice. A moderately-sized lymphoid aggregate and/ or more linear cortical strand with mild lymphocytic infiltrates and compressed tubules were seen in two kidneys from each group of diabetic mice with or without treatment with ND-13 or MCC950. The average mid-cortex capillary glomeruli area in control and diabetic mice showed a minimal difference (34.2 ± 0.18 and 35.2 ± 0.18, respectively) and cell counts in these glomeruli were not significantly different (50.0 ± 8.4 vs 41.8 ± 9.1, respectively).

Assessments of tubular injury and capillary glomerular area in the five largest glomeruli per mouse showed a significant decrease in glomerular capillary area between diabetic mice and normal controls without a significant difference in total glomerular area. Treatment of diabetic mice with ND-13 or MCC950 showed a significant decrease in tubular injury and mean glomerular capillary areas that were comparable to the untreated controls (**Figure 5a-c**).

### 2.5 Patients with diabetic nephropathy have IL-1β release increased

We analysed peripheral mononuclear cells (PBMCs) from patients with diabetes, diabetic nephropathy and healthy donors. We stimulated PBMCs with LPS+ATP, it was confirmed a significant difference between diabetic patients and DN patients in the release IL-1β (52.44±23.50 vs 255.5±83.78; n=11-14, Mann Whitney test, **p<0.005) (**Figure 6a**) suggesting a significant activation of NLRP3 inflammasome in DN. However, TNF-α release did not offer any significant difference (**Figure 6c**). Although ND-13 treatment of PBMCs did not show decreased release of IL-1β or TNF-α in any of the groups, a possible reduction in IL-1β is seen in healthy donor cells treated with ND-13 (**Figure 6b**), suggesting that ND-13 may be more effective in preventing inflammasome activation.

**Figure 6.**
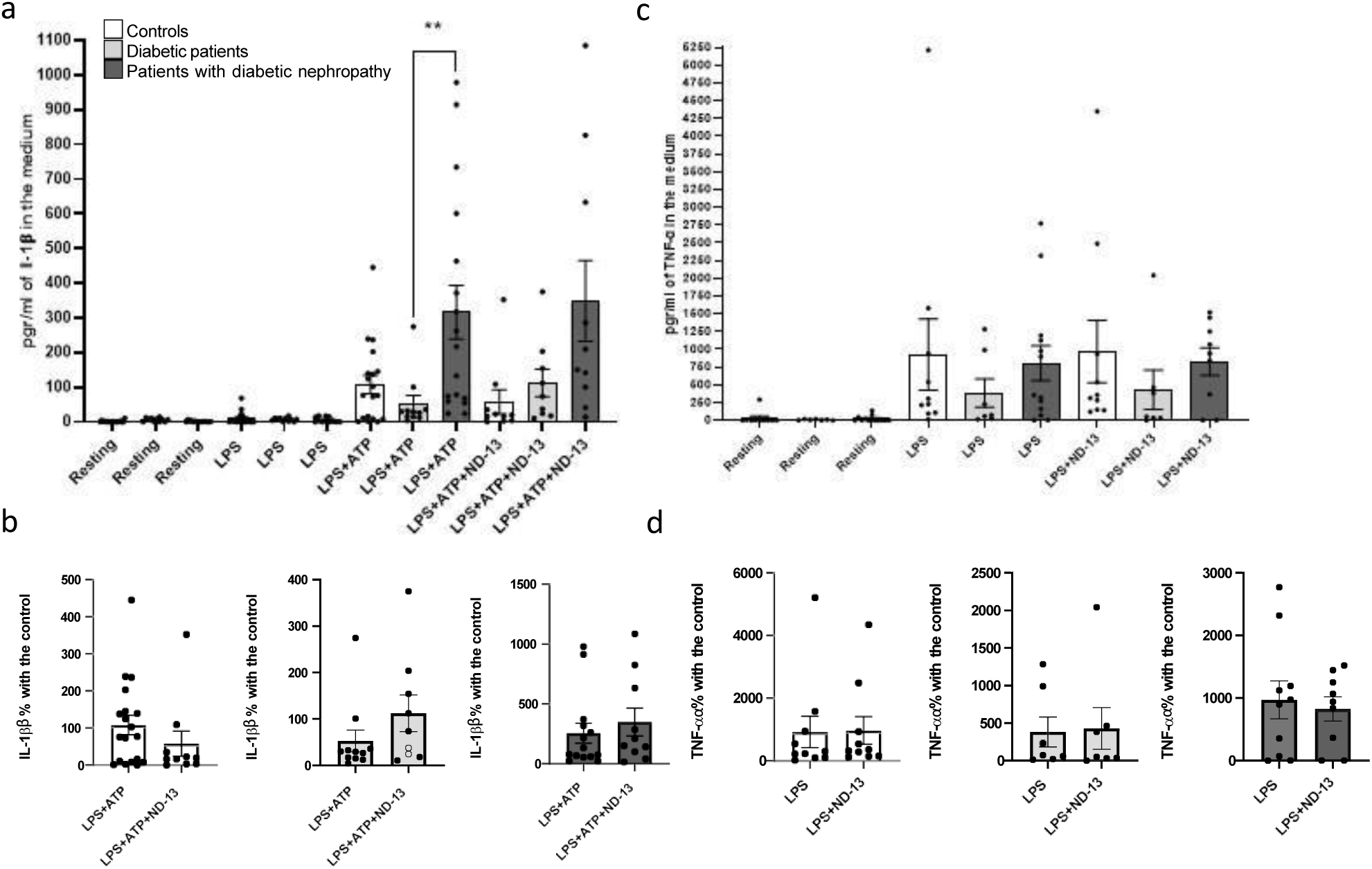
NLRP3 inflammasome and NFκB pathway activation in human. **(a-b)** IL-1β release to the medium, a marker of inflammasome activation, in peripheral blood mononuclear cells (PBMCs) from diabetic patients, patients with diabetic nephropathy and controls. Control n=19, diabetic patients n=11, diabetic nephropathy patients n=17. **(c-d)** TNF-α release to the medium as a marker of NFκB pathway activation via LPS in peripheral blood mononuclear cells from diabetic patients, patients with diabetic nephropathy and controls. Control n=10, diabetic patients n=7, diabetic nephropathy patients n=14. Data is represented as mean ± SEM; Mann Whitney test was used to compare groups in a-d. Significance level is indicated as follow: *\*\*p<0.005*.

## 3. Discussion

This work investigates the association between NLRP3 inflammasome activation with DN and the beneficial effects of the DJ-1 pathway, specifically ND-13, in preventing inflammasome overactivation. This study demonstrates the ability of ND-13 to attenuate inflammasome activation in LPS+ATP activated BMDM. Patients and STZ mice have increased NLRP3 inflammasome activation in PBMCs and peripheral immune cells respectively; this NLRP3 inflammasome overactivation is associated with renal pathogenesis and was normalized in mice by ND-13 treatment and by MCC950, an inhibitor of NLRP3 inflammasome. In addition, N-13 prevents the tubular damages and decrease of the total surface area of all glomerular capillaries induced by STZ and attenuates the expression of interleukins associated with inflammatory response and fibrosis, which are overexpressed in diabetic mice. The data of the present study demonstrate the protective effects of ND-13 to prevent the inflammasome NLRP3 activation in cell culture and in animal models, suggesting that DJ-1-derived compounds may be new strategies of treatment to prevent the development of DN and other renal diseases associated with oxidative stress and inflammation, which may have significant impact in the field.

The inflammasome is a multi-protein complex that plays a crucial role in the innate immune response. NLRP3 inflammasome can be activated by a variety of stimuli, including metabolic stress (reviewed in [37]), endoplasmic reticulum stress [38], or the release of DAMPs [39]. Thus, kidney disease associated to diabetes factors such as hyperglycaemia, uremic toxins, and oxidative stress may activate the NLRP3 inflammasome and trigger the renal damage [40]. Inflammasome activation release of IL-1β and IL-18 induces a potent inflammatory response, leading to the recruitment of immune cells and the amplification of the inflammatory process. In the kidney, this activation in peripheral cells may increases the infiltration of activated cells which may result in tissue damage, fibrosis, and the progression of kidney disease. In fact, factors associated with cells infiltration such as intercellular adhesion molecule 1 (ICAM-1) are associated with renal diseases [41] and with NLRP3 inflammasome activation in diabetic mice [42]. Previous reports in human with chronic kidney disease (CKD) shown that, IL-1β inhibition with canakinumab reduces the cardiovascular event rates among high-risk atherosclerosis patients, without adverse clinical renal events [43]. NLRP3 inflammasome activity is also associated with glomerular injury in diabetic nephropathy patients [44], with increased glomerular NLRP3 expression [45] suggesting the role of NLRP3 in different cells type in the pathogenesis of diabetic cardio-renal diseases. However, there are not studies that determine the inflammasome activation in the immune cells of these patients. In this study for the first time, PBMCs from diabetic patients with and without nephropathy has been isolated and the NLRP3 inflammasome activation has been determined. Our data show a significant increase in inflammasome activation in patients with diabetic nephropathy compare with diabetic patients, suggesting that an increase of inflammasome activation in blood immune cells is associated with renal damage in diabetic patients, interestingly, this different have not been found in healthy control. This may suggest that diabetic patients without morbidity may have a compensatory mechanism that prevents inflammasome activation and kidney damage.

It is important to note that the type of renal cells involved in the inflammatory response and the type of inflammasome are issues that are still not clear to understand the positive feedback activation of inflammation, oxidative stress and fibrosis that results in renal pathogenesis. Protein expression of other inflammasomes, such as NLRC4, has been described in tubular cells and may influence renal tubular epithelial cell injury in DN [46]. Previous studies have shown a significant increase in the expression of IL-1β, IL-18, ASC, NLRP3, caspase 1 in human umbilical vein endothelial cells (HUVECs) in high glucose conditions [40]; and human kidney-2 (HK-2) proximal tubule cells when treated with high glucose (30 mM) exhibit an increased mRNA and protein expression of NLRP3, ASC, caspase-1, active IL-1β and active IL-18 [47]. In addition, podocyte-specific NLRP3 inflammasome activation may promote diabetic kidney disease [48] and high glucose-treated in primary murine podocytes increased NLRP3 expression impaired podocyte autophagy and exacerbated podocyte damage [49]. Therefore, the increased of expression of NLRP3 inflammasome components have been demonstrated in different types of renal cells such as proximal tubular cells and podocytes, however, the inflammasome activation is not associated with the expression of their compounds and the gasdermin activity, their capacity to induce pyroptosis and the systemic immune response is still not clear in tubular cells and podocytes. We propose that NLRP3 inflammasome activation in immune cells may be a key sensor to initiate the inflammatory response in the renal tissue, making the immune cell response in the presence of high glucose particularly important as a triggering inflammatory mechanism in diabetes disease. Other recent studies have investigated the effect of high glucose on inflammasome activation in the presence or absence of air pollution fine particulate matter (PM2.5) in murine BMDMs; high glucose pretreatment enhanced the effects of PM2.5 on NLRP3 inflammasome activation, but no effects of high glucose alone were detected [50]. We are investigated whether high glucose can increase the inflammasome activity in BMDMs stimulated with LPS, ATP and nigericin, and no significant changes in inflammasome activation have been shown to be influenced by glucose concentration, suggesting that NLRP3 inflammasome activation in immune cells may be influenced by multifactorial mechanism involved in the systemic homeostasis which could be affected in diabetic patients.

Renal fibrosis and renal infiltration, both associated with renal damage, were not found in our diabetic mice as shown in **Figure 4**. Previous publications have reported that STZ diabetic-induced mice start to develop renal damage after 10 weeks and reach maximum damage at 18 weeks [51]. One of the main objectives of this study is to determine whether NLRP3 inflammasome activation trigger renal damage in diabetic. The feedback loop of oxidative stress, inflammation and fibrosis associated to the renal pathological conditions (**Figure 7**) makes it difficult to figure out the sensor mechanism that initiated the inflammatory response in DN. Therefore, analyzing the first steps of inflammatory damage in an incipient diabetic nephropathy may be a good condition to determine the original mechanism involved in the inflammatory response in diabetic conditions. In this contest, we observed inflammasome activation in the peritoneal immune cells of these animals and an incipient inflammation in the renal cortex which may demonstrate the important role of NLRP3 in the initial development of renal pathology.

**Figure 7.**
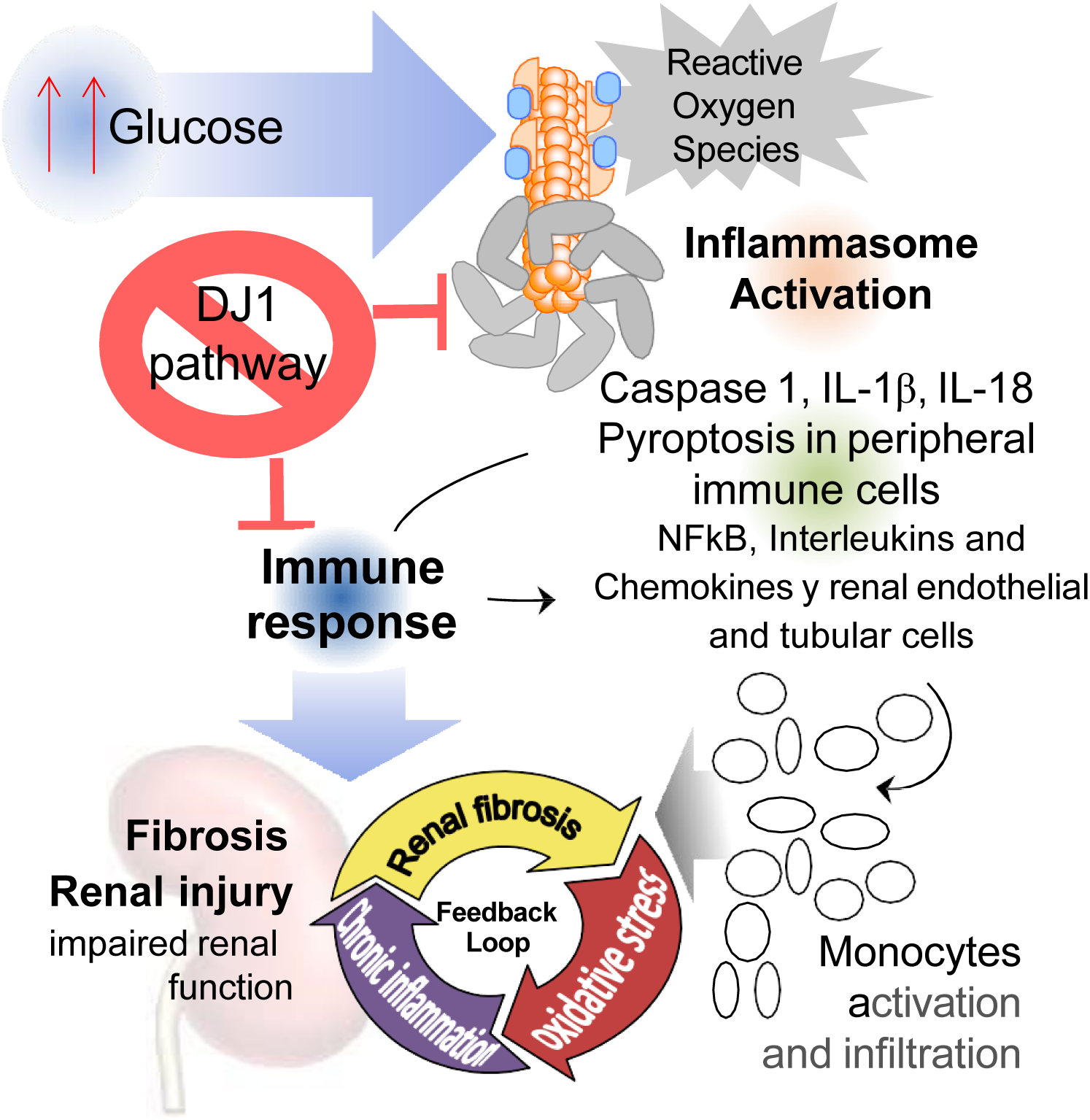
Graphical abstract. High Glucose activates NLRP3 inflammasome and the release of IL-1β and IL-18, which induce the immune responses and the renal infiltration of the immune cells, inducing NFkB translocation, and an increase of ROS production. These stimuli may have a feedback activation loop lending fibrosis, renal injure, and impair renal function. DJ-1 pathway inhibits inflammation, attenuating inflammasome activity preventing renal damage.

The second objective of this study was to ascertain the impact of ND-13 on inflammasome regulation and its anti-inflammatory effects in renal tissue in a diabetic mouse model. Diabetes was induced in C57Bl/6 mice via injection of Streptozotocin (STZ) and treated with ND-13 and MCC950, a NLRP3 inhibitor. The peritoneal immune cells of these mice exhibited an increased production of IL-1β in response to LPS+ATP stimulation, a result that aligns with our findings in humans. Furthermore, ND-13 and MCC950 restored IL-1β production (**Figure 3d**) and renal expression of inflammatory and fibrotic factors induced by STZ in diabetic mice (**Figure 5d-e**), indicating that the DJ-1 pathway may be a promising avenue for attenuating the renal inflammatory response in diabetes. The anti-inflammatory effect of ND-13 in renal diseases has been demonstrated in previous studies [26], however, this is the first time that its effects on inflammasome NLRP3 activation have been shown.

The total length of the capillaries in the glomerulus is associated with the capacity of the filtration slit surface area, and a reduction in the glomerular capillary surface area affects the filtration surface area and renal function. [52]. Thus, atrophy and obliteration of glomerular capillaries compromise glomerular function even in the absence of podocyte lesions. [52]. Therefore, the observed decreased glomerular capillary area may be the first step for the glomerulonephritis induced by my diabetes and ND-13 and MCC950 prevent it, demonstrating their ability to protect renal function. Moreover, tubular damage has also been observed in diabetic animals, which may compromise tubular reabsorption [53], with significant improvement induced by our experimental treatments (**Figure 5b**). The tubular damage in diabetes and the role of inflammation and oxidative stress on this effect is well known [54]. Thus, our data demonstrate the possible key role of inflammasome NLRP3 in the incipient glomerulus and tubular damage in diabetic conditions and the preventive effect of ND-13 on renal function.

P2X7 is a type of purinergic receptor that plays a pivotal role in the immune system and the process of inflammation. It is an ATP-gated ion channel, which means that it opens in response to the binding of ATP, thus allowing the flow of ions such as calcium and potassium across the cell membrane. This ion flux can result in several downstream effects, including the activation of the NLRP3 inflammasome. The results demonstrate that pretreatment of ND-13 inhibits the expression of P2x7 in renal cortex of mouse, which aligns with the observed trend in P2X7 expression in peritoneal cells (**Figures 3e**). The effects of ND-13 in BMDM are evident in cells treated with ATP, but not in cells treated with Nigericin, which affects the membrane permeability and reduces cytoplasmic potassium concentration via P2X7 independent mechanism. It may therefore be surmised that the effects of ND-13 are mediated by a reduction in P2X7 expression. In diabetes, particularly Type 2 diabetes, P2X7 receptor has been identified as a key contributor to the chronic inflammatory state that characterizes the disease. The activation of the receptor on immune cells results in the release of pro-inflammatory cytokines, which have the potential to interfere with insulin signaling pathways in tissues such as adipose tissue, liver, and muscle, thereby contributing to insulin resistance [55]. Additionally, P2X7 has been demonstrated to induce β-cell dysfunction, leading to β-cell stress and apoptosis [56]. Furthermore, it has mediated activation of the inflammasome and subsequent cytokine release in diabetes [57].Therefore, the P2X7 receptor plays a substantial role in the inflammatory processes associated with diabetes. It can be hypothesized that the ND-13 effects may be mediated, at least in part, by the downregulation of P2X7 expression. It is important to note that the expression of P2X7 is not exclusive of immune cells. Previous studies have associated an increase of P2X7 expression in podocytes in diabetes, which have been show to inhibit podocyte autophagy and exacerbate podocyte damage, thereby promoting the onset of diabetic nephropathy [58]. The impact of ND-13 on P2X7, NLRP3 and PYCARD expression is more pronounced in renal cortex that in contrast of the peritoneal cell, where a trend is observable, but a significant effect has not been identified. This suggests that the effects of ND-13 may be mediated in renal cell such as podocyte and tubular, and that it effects are not solely dependent on immune cells. Nevertheless, the impact of NLRP3 inflammasome activation on renal cells remains unclear. Additionally, the expression of IL-18 and IL-1β remains unaltered in the renal cortex of these animals, in contrast to other inflammatory cytokines such as IL-6 and TNF-α. This aligns with our previous findings [26], as both cytokines are independent of inflammasome activation. This suggests that the role of the inflammasome in triggering the inflammatory response may be mediated by the effects of other cytokines in other cell types within the tissue, contributing to the activation of further inflammation and the perpetuation of a pathogenic inflammatory vicious cycle.

Similarly, other inflammatory pro-fibrotic cytokines were increased in diabetic mouse. The development of renal fibrosis is influenced by the accumulation of extracellular matrix components, including collagen [59]. Macrophages are a primary source of transforming growth factor beta (TGF-β) in fibrotic organs [60, 61], and recruitment of T- and B-lymphocytes to the site of injury further facilitates secretion of fibrogenic cytokines [62, 63]. TGF-β is also a potent chemoattractant, facilitating the recruitment of inflammatory cells [64], and thereby the expansion of the inflammatory process. TGF-β have been consider by several authors as a master regulator of DN [65]. Our findings demonstrate that the expression of collagen and TGF-β is elevated in STZ-induced diabetes and ND-13 models, and that MCC950 can attenuate these expressions, as previously demonstrated in renal disease models [26].

The transcription factor Nrf2 (nuclear factor erythroid 2-related factor 2) is a master regulator of the expression of several antioxidant genes. Previously published research has indicated that GSK3 plays a role in regulating the proteasomal degradation of NRF2. Consequently, GSK3 inhibition may be an effective strategy for protecting cells from oxidative stress and increasing NRF2 activity [66]. As previously demonstrated in our publication, renal DJ-1 increases Nrf2 stability and prevents its proteasome degradation [20]. Additionally, it has been reported that DJ-1’s cardioprotective effect in diabetic rats is mediated by PTEN inhibition and the activation of Nrf2/HO-1 [67]. PTEN regulates GSK3 via the AKT pathway; thus, it can be postulated that DJ1 activity may increase GSK3 inhibition via the inhibition of PTEN [68]. Based on the aforementioned evidence, we sought to ascertain whether ND-13 exerts an effect on GSK3 inhibition. However, no discernible alterations in GSK3 phosphorylation were observed in cells and mice treated with ND-13 (**Supplementary figure 2**). Furthermore, as anticipated, Nrf2 activity was observed to be elevated in diabetic mice, while the expression of Nrf2 target genes, including HMOX1, NQO1 and GTA1, was found to be diminished in mice treated with ND-13. It has not been possible to confirm whether NRF2 is involved in the protective effect of ND-13. This may be due to the fact that the effects occur at a specific point in time, making it difficult to determine whether Nrf2 has been activated previously. Conversely, a recent report indicates that DJ-1 mitigates the glycation of mitochondrial complex I and complex III in the post-ischemic heart [66], however, it remains to be seen whether this or other effect are associated with ND-13 treatment in our animals.

Activation of the NLRP3 inflammasome plays may be a key factor in the inflammatory and metabolic dysfunction observed in diabetes. This study determines for the first time the human NLRP3 activation in PBMCs and shows interesting differences between diabetic and diabetic nephropathy patients, suggesting the important role of activation of peripheral immune cells in the renal damage associated with diabetes may serve as a potential marker for early disease prevention. Understanding the mechanisms underlying NLRP3 activation and its impact on insulin resistance may inform the development of targeted therapies to alleviate the inflammatory component of diabetes and improve patient outcomes. In addition, we highlight the DJ-1 pathway, and particularly our peptide ND-13, as a novel potential pharmacological approach to attenuate renal fibrosis and inflammation and prevent renal damage in DN.

## 4. Material and Methods

This research complies with all relevant ethical regulations and the study protocol for patients was approved by the ethical committee of the University Clinical Hospital Virgen de la Arrixaca (Murcia, Spain). All animal protocols were conducted in accordance with the Guide for the Care and Use of Laboratory Animals and were approved by the ethical committee of animal experimentation of the University of Murcia. Human’s volunteers included in this study gave written informed consent.

### Reagents

Different reagents used in this study were: ATP (A2383, Sigma-Aldrich), Nigericin sodium salt sodium (N7143, Sigma-Aldrich), MCC950 (CP-456773), L-glutamine (Cat. No: SH30034.01, Hyclone laboratories), Foetal Bovine Serum (Invitrogen Life Technologies), penicillin+streptomycin (P/S) and LPS ultrapure (tlrl-3pelps, Invivogen). Ultrapure water filter through Milli-Q system (Millipore Corp).

### Mouse bone marrow-derived macrophages (BMDMs)

BMDMs were differentiated from precursors obtained from C57BL/6 (wild type, WT) mice between 8 and 10 weeks of age bred under SPF conditions. Bone marrows were obtained from femurs and tibias of mice euthanized by CO_2_ inhalation and resuspended in DMEM High Glucose (L0102, Biowest) with 10% FBS, 100 U/ml penicillin, and 100µg/ml streptomycin, supplemented with 20% conditioned medium from L929 cells as previously described [69]. After 6 days, resulting BMDMs were detached with cold PBS 1X, counted, and plated at a concentration of 500.000 cells per well in plates of 24 wells and in a final volume of 500μl. Plates were incubated at least 3 hours at 37° and 5% CO_2_ before stimulation of cells to ensure that macrophages adhered well to wells.

• Stimulation assay: ND13 treatment

After previous incubation, medium was removed from all wells and 500μl of Opti-MEM™ (51985034, Gibco) per well was added. Appropriate wells were treated with ND-13 (4uM) and plates were incubated overnight at 37° and 5% CO_2_ to ensure the action of the peptide. LPS ultrapure (100ng/ml) was added, and plates were incubated for 3h at 37°C and 5% CO_2_. Medium was collected after 3h, centrifuged at 13200 rpm for 30 seconds at 4°C, transferred to a new tube and stored at -80°C for future ELISA detection of TNF-α. Then, 300μl of physiological E-total buffer (pH 7.45, 147mM NaCl, 10mM HEPES, 13mM D-glucose, 2mM KCl, 2 mM CaCl_2_, 1mM MgCl_2_) and nigericin (10μM) were added and an incubation of 10 minutes was done. After 10 minutes, ATP (3mM) was added, and plates were incubated 20 minutes more. After this time, supernatants were collected, centrifuged at 13200 rpm for 30 seconds at 4°C, transferred to a new tube and stored at -80°C for their future analysis.

• GSK3/P-GSK3 assay

BMDM were plated at a concentration of 2.000.000 per well in plates of 6 wells. After at least 3 hours, medium was removed from all wells and 500μl of Opti-MEM™ (51985034, Gibco) per well was added. Appropriate wells were treated with ND-13 (4uM) and plates were incubated overnight at 37° and 5% CO_2_ to ensure the action of the peptide. Cells from each well were collected using cold lysis buffer (50mMTris-HCl pH 8,0, 150mMNaCl, 2% Triton X-100) supplemented with 100μl/ml of protease inhibitor mixture (P8340 Sigma-Aldrich). Supernatant was collected and incubated in ice for 30 minutes, then it was centrifugated at 13200 rpm for 10 minutes at 4 °C and transferred to a new tube and stored at -80°C for their future analysis.

#### BMDM in high glucose vs low glucose

BMDMs were differentiated from precursors obtained from C57BL/6 (wild type, WT) mice between 8 and 10 weeks of age bred under Specific-pathogen-free (SPF) conditions. Bone marrows were obtained from femurs and tibias of mice euthanized by CO_2_ inhalation and resuspended in DMEM low glucose (D5546, Sigma-Aldrich) with 15% FBS, 1% penicillin/streptomycin, 584mg/L of L-glutamine, 25 ng/ml of Recombinant Murine M-CSF (315-02, Peprotech) and, only in the case of BMDM cultivate in hight glucose, 4,5 g/L of extra glucose was added to the medium. After 6 days, resulting BMDMs were detached with cold PBS 1X, counted, plated and incubated as previously described but in DMEM low glucose with 20% FBS, 1% penicillin/streptomycin, 584mg/L of L-glutamine and, only in the case of BMDM cultivate in hight glucose, 4,5 g/L of glucose. After overnight incubation, medium was removed from all wells and 500μl of Opti-MEM™ (51985034, Gibco) per well was added. LPS ultrapure (100ng/ml) was added, and plates were incubated for 3h at 37°C and 5% CO_2_. Nigericin (10μM) and MCC950 (10μM) were added, and an incubation of 10 minutes was done. Then, ATP (3mM) was added, and plates were incubated 20 minutes more. Finally, supernatants were collected, centrifuged at 13200 rpm for 30 seconds at 4°C, transferred to a new tube and stored at -80°C for their future analysis.

#### Cell death assay

Cells used as positive control were scraped using cold lysis buffer (50mMTris-HCl pH 8,0, 150mMNaCl, 2% Triton X-100) supplemented with 100 μl/ml of protease inhibitor mixture (P8340 Sigma-Aldrich). Supernatant was collected and incubated in ice for 30 minutes, then it was centrifugated at 13200 rpm for 10 minutes at 4 °C, transferred to a new tube and stored at -80°C for their future analysis. A colorimetric assay was performed to quantify cell death and cell lysis, based on the measurement of lactate dehydrogenase (LDH) activity released from the cytosol of damaged cells into the supernatant (ROCHE Cytotoxicity Detection Kit (LDH)). Plates were read in a Synergy Mx (BioTek) plate reader at 492 nm and corrected at 620 nm.

#### ELISA assays

Cell-free supernatants were collected, and the following ELISA kits were used according to manufacturer instructions: mouse IL-1β (88-7013-88 or 88-7013A-88, Invitrogen), mouse TNF-α (88-7324-88, Invitrogen), human IL-1β (BMS224INST, Invitrogen) and human TNF-α (DY210, R&D Systems). ELISA were read in a Synergy Mx (BioTek) plate reader at 450 nm and corrected at 570 or 620 nm.

#### Animals and treatments

All animal protocols were conducted in accordance with the Guide for the Care and Use of Laboratory Animals and were approved by the ethical committee of animal experimentation of the University of Murcia (reference CEEA: **646/2020**) and the Animal Health Service of the General Directorate of Fishing and Farming of the Council of Murcia (reference **A13230110**). C57BL/6 mice (WT, wild-type, RRID: IMSR_JAX:000664) were obtained from the Jackson Laboratories. Mice were bred in specific pathogen-free conditions with a 12:12 h light-dark cycle and used in accordance with the Spanish national (Royal Decree 1201/2005 and Law 32/2007) and EU (86/609/EEC and 2010/63/EU) legislation. For all experiments, diabetes was induced in 8-9 weeks male mice via intraperitoneal injection of 50 mg/kg of streptozotocin (STZ) for 5 days using citrate buffer pH 4.5 as diluent while control mice were intraperitoneal injected with citrate buffer pH 4.5. Diabetes mice were treated with ND-13 (3 mg/kg) or MCC950 (10mg/kg) 3 days a week for 10 weeks. Weight was measure before and after treatment, whereas kidneys, peritoneal macrophages and final point blood were harvested after treatment and sacrifice of mice with CO_2_. One of the kidneys was snap-frozen in liquid nitrogen for qPCR, while the other was placed in formalin for histological studies. Glucose concentration was measured in blood from a slight cut in the tail of animals by a glucometer except at the end point where blood was obtained directly from the heart of each mouse.

#### Mouse peritoneal macrophages assay

Peritoneal macrophages were obtained by flushing the peritoneal cavity with 5 ml ice-cold PBS. Cells were then centrifuged at 1500 rpm for 5 minutes, counted, resuspended in complete RPMI medium (RPMI R0883, Sigma-Aldrich with 10% FBS, 1% L-Glutamine y 1% P/S) in the appropriate amount to remain at 1mill/ml of cells and plated in 24-well plates at a concentration of 250.000 cells per well, to a final volume of 500μl of complete RPMI medium. Plates were incubated overnight at 37° and 5% CO_2_ to ensure that macrophages adhered well to the wells.

#### Stimulus preparation

After overnight incubation, we remove medium from all wells and add 500μl of complete RPMI medium to the wells that do not contain any stimulation (control wells) and 500μl of complete RPMI medium+LPS ultrapure stock (100ng/ml) to the remaining wells that were going to be stimulated or not with ATP. We leave plates for 3h at 37°C and 5% CO_2_, then we washed with 500μl of ET-buffer (147 mM NaCl, 10 mM HEPES, 13 mM D-glucose, 2 mM KCl, 2 mM CaCl_2_, and 1 mM MgCl_2_; pH 7.4) per well twice, added 500μl of ET-buffer in all wells and ATP (3mM) in the appropriate wells. After 30 min, supernatants were collected, centrifuged at 13200 rpm for 30 seconds at 4°C, transferred to a new tube and stored at -80°C for their future analysis. Cells were lysed with 250μl per well of RLT + 1% beta-mercaptoethanol a scraper and transfer to new tubes at -80°C.

#### Quantitative reverse transcriptase-PCR analysis

Total RNA was purified and isolated from renal cortex or peritoneal macrophages from mice using the RNeasy kit (74104, Qiagen) following manufacturer instructions and quantified in a NanoDrop 2000 (Thermo Fisher). RNA was reverse transcripted using the iScriptTM cDNA Synthesis kit (1708891, BioRad) according to manufacturer instructions. Quantitative PCR was done in an iQTM 5 Real Time PCR System (BioRad) with SYBR Green mix (Takara) and predesigned KiCqStart primers for mouse *Il1b, Il6, Tnfa, Il18, Col I, Col IV, Tgfb, Pycard, Nlrp3, Nek7, P2x7, Casp1, Nqo1, Gsta1* and *Hmox* (Sigma-Aldrich). Gene expression was normalized with β-actine as endogenous control.

#### Histology, Fibrosis Quantification and Quantification of Immune Cell Infiltration in the Kidney

Kidney samples were fixed in 4% buffered formalin (Panreac Quimica, Barcelona, Spain) for 12 hours, processed, paraffin embedded and 4 µm-thick sectioned. For histopathologic examination, kidney sections were stained with a standard haematoxylin and eosin stain. Additionally, to determine the degree of fibrosis, a Masso’s trichrome stain was performed in all samples by using a commercial kit (04-010802, Bio-Optica, Nmilano, Italy) following the supplie’s recommendations.

To determine the presence of T cells and macrophage infiltration within kidney tissue, an indirect based-polymer colorimetric immunohistochemistry was carried out in all samples. Briefly, after deparaffination and rehydration, sections were demasking antigen treated by using a commercial solution (Dako target retrieval solution, Agilent Technologies, Madrid, Spain). After peroxidase-blocking, sections were overnight incubated with an specific primary antibodies (polyclonal rabbit anti-CD3, A0452, Dako-Agilent, dilution 1:500, and monoclonal rat anti-F4/80, AB6640, Abcam, Oxford, UK, dilution 1:250) and a secondary-labelled biotin-free HRP labelled polymer (Vector ImmPress anti-rabbitand anti-rat, Vector Labs., Burlingame, USA) for 20 min at room temperature. Finally, the immunoreaction was revealed by incubating the slides with 3-3’diaminobencidine (DAB). Positive immunoreaction was evidenced by a dark-brown precipitated with membrane pattern.

For histopathologic examination, all slides were digitalized by using a brightfield high resolution slide scanner (Pannoramic MIDI II, 3D Histech, Budapest, Hungary). The cortical area of each fμll kidney scan was outlined and the percentage of blue fibrotic deposition within the outlined area was quantified using MetaMorph software (Molecμlar Devices LLC., San Jose, CA, USA). The average percentage fibrotic area for each experimental group was calculated and then normalized to the control group. Data are presented as the average percentage area that stained positively per experimental group. Quantification of renal macrophages (F4/80+ cells) and T-lymphocyte (CD3+ cells) infiltration was performed by taking 10 cortical images from each animal at 400× magnification. The percentage of cortical area stained positive for F4/80 or CD3+ was quantified in each image using MetaMorph software and the average expression per animal was calculated. The data are reported as the average percentage area that stained positively for F4/80 or CD3+ per experimental group.

#### Histologic Methods for tubular and glomerular analysis

Formalin-fixed paraffin-embedded tissues were sectioned from one kidney per mouse at 3-4 microns and routinely stained with H&E. Kidney pathology was evaluated in 8 normal controls and 10 mice treated with STZ. Slide sections were used for histologic analyses after whole slide scanning (Aperio CS2) and viewed on SlideViewer (3D Histotech). Renal pathology was assessed for evidence of inflammatory infiltrates and injury to vasculature, glomeruli and tubules. The number of tubules showing injury was assessed in 10 areas of cortical tissue (at 10X or 2.3 mm) per mouse and is reported as the average number of affected tubules/2.3 mm cortex for mice in each category. Used the capillaries of glomerular tuft (not including Bowman’s space) for the cell counts because that is where any proliferation of cells would be seen at first in most cases. Five of these capillaries per mouse were selected from the mid-cortex on the basis of a roughly round contour and evident hilus. The areas of the 5 mid-cortex capillary glomeruli were calculated from their diameters and cell counts were determined and are reported as averages for the mice in each category. The five largest glomeruli per mouse were also evaluated to compare the capillary tuft area and the renal corpuscle area (tuft plus Bowman’s space) as measured by the annotation tool of their circumferences circumferences in SlideViewer [70]. The histologic features that showed significant differences between the control and STZ-treated mice were further assessed in 7 mice treated with ND-13 and 8 mice treated with MCC950, with significance differences determined by Students T-test.

#### Western blot

Kidneys samples were grinded with a tissue homogeniser until the sample was as homogeneous as possible. They were centrifugated at 1300rpm for 15 minutes at 4°C and supernatants collected. The supernatants were quantified using the Bradford assay (Sigma-Aldrich) and the results were read on a Synergy Mx plate reader (BioTek) at 595nm. Samples in which Synaptopodin and Desmin proteins were measured were diluted at a proportion of 1:10 in water. Supernatants were denatured with Laemmli buffer (Sigma-Aldrich) and heated at 95°C for 5 minutes. 40μg of concentrated supernatants were resolved on 12% criterion polyacrylamide gels and transferred to nitrocellulose membranes (Biorad) by electroblotting. Membranes were incubated with 100 rpm shaking for 1 hour and at room temperature with 5% milk in 1x TBS-T+0.05% Tween. Membranes were then hybridised overnight at 4°C with the following primary antibodies: Synaptopodin (sc-515842, Mouse; 1:1000), Desmin (sc-23879, Mouse; 1:1000), GSK3 (sc-7291, Mouse; 1:1000) and P-GSK3 (1679331S, Rabbit; 1: 1000) and the following day with the appropriate secondary antibodies: Synaptopodin (NA931, Mouse; 1:5000), Desmin (NA931, Mouse; 1:5000), GSK3 (NA931, Mouse; 1:5000) and P-GSK3 (NA9340, Rabbit; 1:5000). Membranes were developed using ECL Plus reagents (Cytiva) on a ChemiDoc imaging system (BioRad). Membranes were then incubated with horseradish peroxidase-conjugated (HRP)-anti-β-actin HRP-bactin (sc-47778, 1:10,000) secondary antibody for 30minutes at room temperature. Band intensities were quantified using Image J software.

#### Human samples

Data and samples were stored in the Region of Murcia Biobank, registered on the National Registry of Biobanks (BIOBANC-MUR B.0000859), and were used following standard operating procedures. Gender was not considered in this study. This research complies with all relevant ethical regulations and the study protocol for patients was approved by the ethical committee of the University Clinical Hospital Virgen de la Arrixaca (Murcia, Spain). Three groups of patients had be included: health no diabetic patients, diabetic without other complications and diabetic with nephropathy. All of them must had signatured an informed consent. Whole blood samples were collected in EDTA anticoagulated tubes from adults aged 18 to 65 years and classified according to the **inclusion and exclusion criteria\*** into healthy donors (n=19), patients with diabetes (n=11) and patients with diabetic nephropathy (n=14). Samples were all used to obtain the results of this study. Human peripheral blood mononuclear cells (PBMCs) were isolated using Ficoll Histopaque-1077 (Sigma-Aldrich). Then, resuspended in Opti-MEM™ and plated at a concentration of 500.000 cells per well in plates of 24 wells. ND-13 was added to some wells and plates incubated overnight at 37° and 5% CO_2_. Next day, cells were stimulated with LPS (2μg/ml) for 3h and then stimulated with ATP (3mM) for 20 minutes. After this time, supernatants were collected, centrifuged at 13200 rpm for 30 seconds at 4°C, transferred to a new tube and stored at -80°C for their future analysis.

### *Inclusion and exclusion criteria

-Healthy donors must had a basal glycemia under 100 mgr/dL and a HbA1c under 5,7%. Not have had an anti-inflammatory treatment in the last month and not have diabetes, cardiovascular or renal diseases or other metabolic diseases.

-Patients with diabetes must had diabetes type 2 (basal glycemia equal to 110 mgr/dL or higher and a HbA1c equal to 5,7% or higher) and a glomerular filtration equal to 60 ml/min/1.73 m2 or higher. Not have had an anti-inflammatory treatment in the last month and not have diabetes type I.

-Patients with diabetic nephropathy must had diabetes type 2 (basal glycemia equal to 110 mgr/dL or higher and a HbA1c equal to 5,7% or higher) and have been diagnosed with nephropathy by the Endocrinology Section (glomerular filtration between 15 and 60 ml/min/1.73 m^2^). They also must not have taken an anti-inflammatory treatment in the last month or have diabetes type I.

### Statistics analysis

All data are represented as mean values and errors bars represent standard error (SEM). Normality was determined with D’Angostino and Pearson omnibus K2 normality test or Saphiro-Wilk normality test, followed by the corresponding parametric or non-parametric test. P value < 0.05 was considered statistically significant. Statistical analyses were performed using GraphPad Prism 9.5.1 for Windows (GraphPad Software, San Diego, California USA).

## Abbreviations

AU: Arbitrary unit
BP: Blood pressure
CKD: Chronic kidney disease
CO_2_: Carbon dioxide
Col IV: Collagen IV
Col1a1: Collagen 1a1
CRP: c-reactive protein
DAMPs: damage-associated molecular patterns
DN: Diabetic nephropathy
BMDM: Bone marrow derived macrophages
FPG: Fasting plasma glucose
GSK3: Glycogen synthase kinase 3
HAMPs: homeostasis-altering molecular processes
HbA1c: Glycosylated haemoglobin
HK-2: Human kidney-2
HMOX1: Heme-Oxygenase-I
HUVECs: Human umbilical vein endothelial cells
ICAM-1: Intercellular adhesion molecule 1
LDH: Lactate dehydrogenase
LPS: Lipopolysaccharide
Nrf2: Nuclear factor erythroid 2-related factor 2
Nrf2: Nuclear factor erythroid 2-related factor 2
PAMPs: pathogen-associated molecular patterns
PBMCs: Peripheral mononuclear cells
PD: Parkinson’s disease
P-GSK3: Phosphoglucose synthase kinase 3
ROS: Reactive oxygen species
SPF: Specific-pathogen-free
STZ: Streptozotocin
TLR: Toll-like receptor
UUO: Unilateral ureter obstruction

## Disclosures

SC and PP are co-founders of Viva in vitro diagnostics SL but declare that the research was conducted in the absence of any commercial or financial relationships that could be construed as a potential conflict of interest. The author Santiago Cuevas declare that hold a provisional patent in the United States on the effect of ND-13 in renal disorders.

## Funding

SC was funded by Fundación Séneca 21921/PI/22, Instituto de Salud Carlos III PI22/00129 co-funded by the European Union. Funding sources provided financial support but were not involved in study design, collection, analysis and interpretation of data.

## Acknowledgments

The authors thank the pathology and genomic platforms of the Murcia Institute of Biosanitary Research of the Region of Murcia (IMIB) for their services and the technical assistance, and IMIB-Arrixaca Biobank to store and administrate human samples. Besides, the Histology Services at the George Washington University Research Pathology Core Lab for their expertise.

## Author Contributions

SC and PP participated in the design of the study and in the discussion and interpretation of the data. IQ performed part of the in vitro experiments. PL to study the morphological damage induced in STZ mice. CM and YZ performed the staining data in STZ mice. MJCH, SC, CM, ADS, DA, and LH performed the human experiments in PMBCs and analyzed the data. DA helps with technical issues during the experimental process. FR, JBC, and LAR are responsible for enrollment and collection of blood samples from patients in the study. MJCH and SC performed the figures, the literature search and drafted the manuscript.

## Data Availability Statement

All data, both clinical and laboratory, will be published following FAIR (Findable, Accessible, Interoperable and Reusable) principles. This will facilitate the results of this study is easily reproducible, and that all the information generated (that does not affect sensitive data) can be easily found and reused in other studies. The human data collection will be carried out in electronic format through a Web platform (SemanticCRF) developed by the Biomedical Informatics and Bioinformatics Platform of IMIB and included in (https://dataverse.harvard.edu/).

## Conflicts of Interest

SC is co-founders of Viva in vitro diagnostics SL but declare that the research was conducted in the absence of any commercial or financial relationships that could be construed as a potential conflict of interest. The author Santiago Cuevas declare that hold a provisional patent in the United States on the effect of ND-13 in renal disorders

---

**Supplementary Figure 1.**
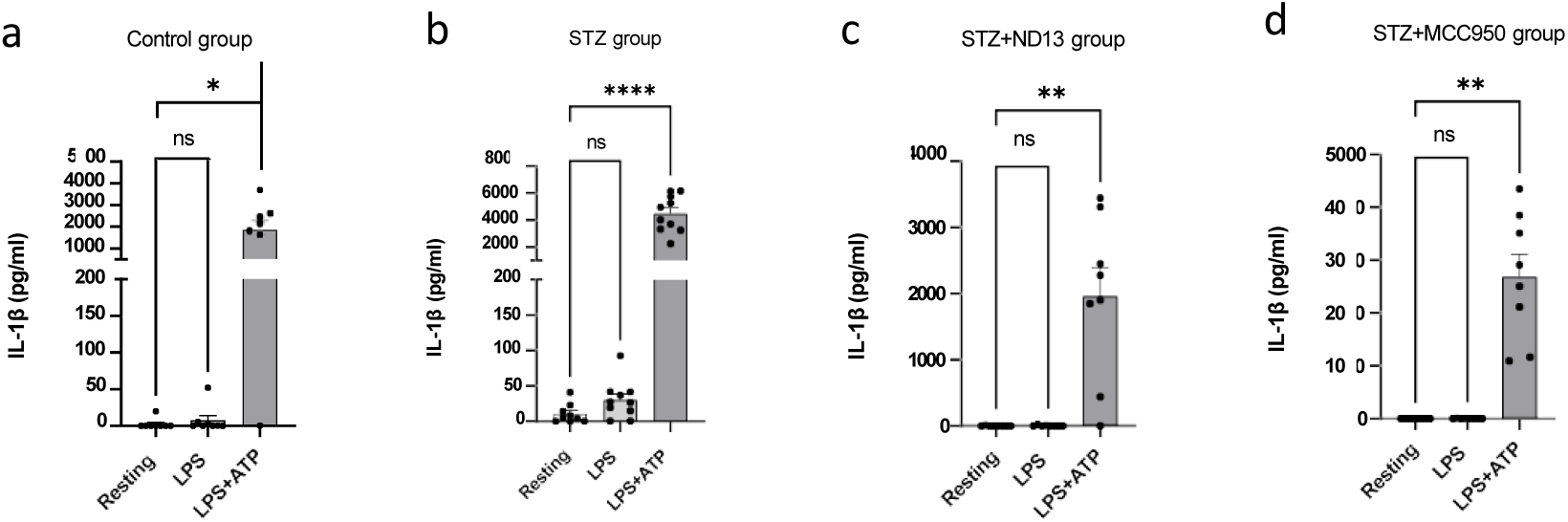

**Supplementary Figure 2.**
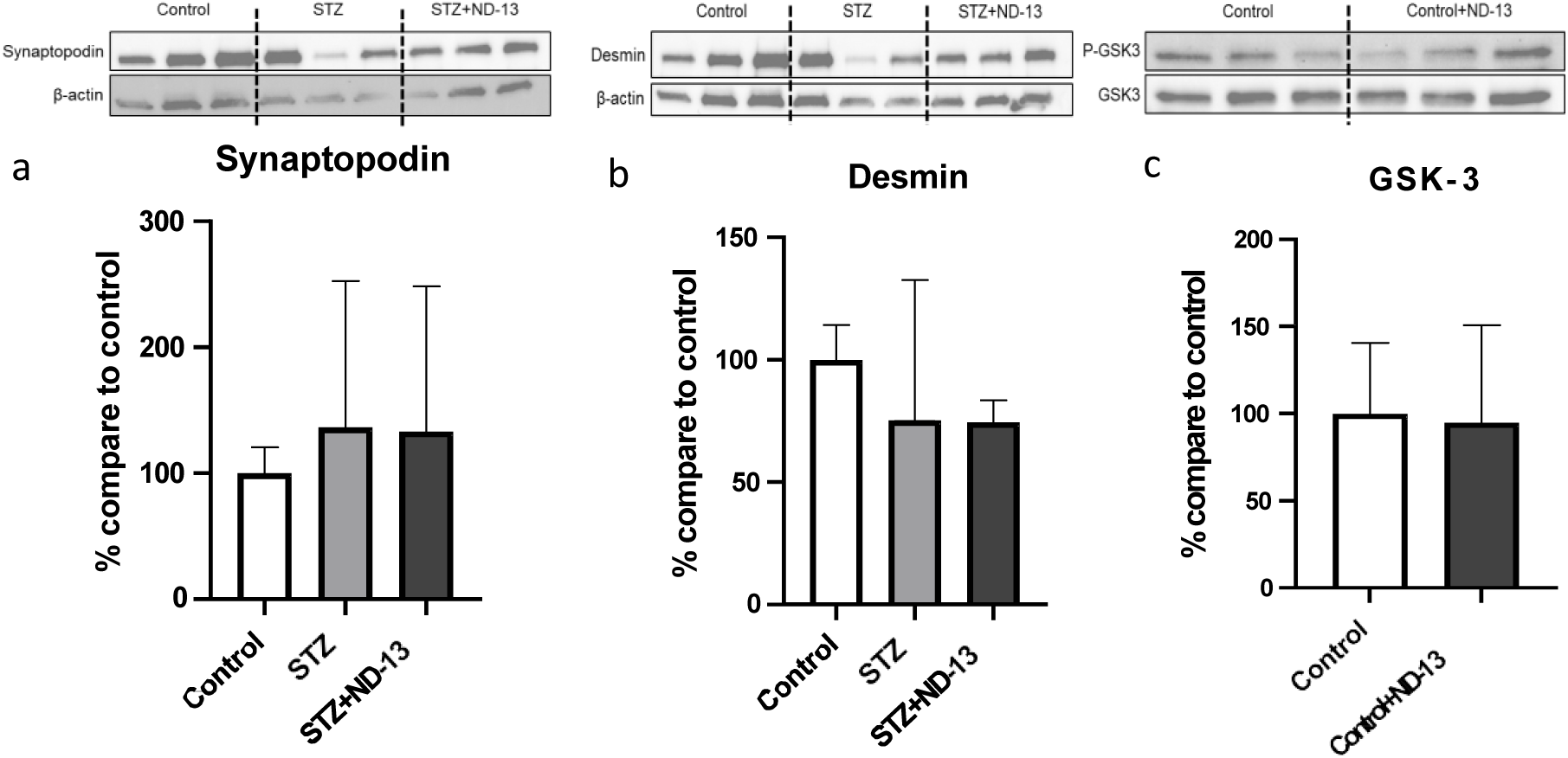

**Supplementary Figure 3.**
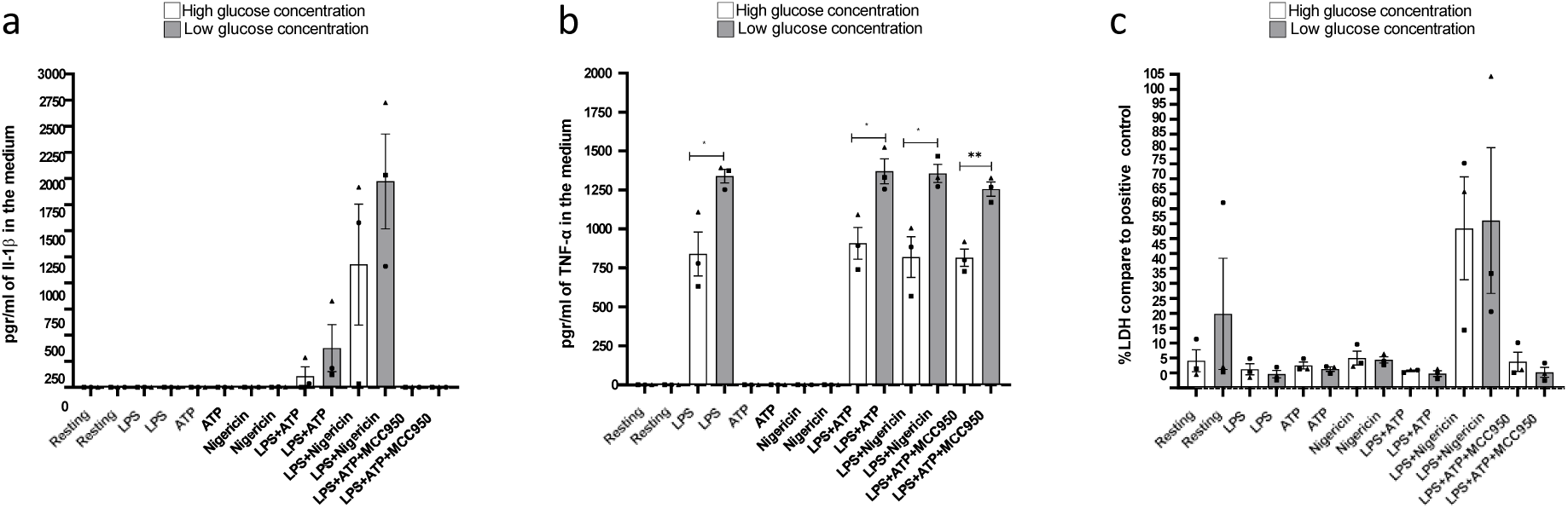

## Notes

https://dataverse.harvard.edu/

## References

1. de Boer, I.H., et al., Diabetes Management in Chronic Kidney Disease: A Consensus Report by the American Diabetes Association (ADA) and Kidney Disease: Improving Global Outcomes (KDIGO). Diabetes Care, 2022. 45(12): p. 3075–3090.

2. Ruilope, L.M., et al., Prevention of cardiorenal damage: importance of albuminuria. Eur Heart J, 2023. 44(13): p. 1112–1123.

3. Giacco, F. and M. Brownlee, Oxidative stress and diabetic complications. Circ Res, 2010. 107(9): p. 1058–70.

4. Higgins, G.C. and M.T. Coughlan, Mitochondrial dysfunction and mitophagy: the beginning and end to diabetic nephropathy? Br J Pharmacol, 2014. 171(8): p. 1917–42.

5. Navarro-Gonzalez, J.F. and C. Mora-Fernandez, The role of inflammatory cytokines in diabetic nephropathy. J Am Soc Nephrol, 2008. 19(3): p. 433–42.

6. Rayego-Mateos, S., R. Goldschmeding, and M. Ruiz-Ortega, Inflammatory and Fibrotic Mediators in Renal Diseases. Mediators Inflamm, 2019. 2019: p. 7025251.

7. Meng, X.M., D.J. Nikolic-Paterson, and H.Y. Lan, Inflammatory processes in renal fibrosis. Nat Rev Nephrol, 2014. 10(9): p. 493–503.

8. Fan, J., et al., Roles of Inflammasomes in Inflammatory Kidney Diseases. Mediators Inflamm, 2019. 2019: p. 2923072.

9. Caruso, R., et al., NOD1 and NOD2: signaling, host defense, and inflammatory disease. Immunity, 2014. 41(6): p. 898–908.

10. Hou, L., et al., NLRP3/ASC-mediated alveolar macrophage pyroptosis enhances HMGB1 secretion in acute lung injury induced by cardiopulmonary bypass. Lab Invest, 2018. 98(8): p. 1052–1064.

11. Broz, P., P. Pelegrin, and F. Shao, The gasdermins, a protein family executing cell death and inflammation. Nat Rev Immunol, 2020. 20(3): p. 143–157.

12. Tartey, S. and T.D. Kanneganti, Inflammasomes in the pathophysiology of autoinflammatory syndromes. J Leukoc Biol, 2020. 107(3): p. 379–391.

13. Kim, Y.G., et al., The Role of Inflammasome-Dependent and Inflammasome-Independent NLRP3 in the Kidney. Cells, 2019. 8(11).

14. Cuevas, S. and P. Pelegrin, Pyroptosis and Redox Balance in Kidney Diseases. Antioxid Redox Signal, 2021.

15. Li, D., et al., Effects of anti-inflammatory therapies on glycemic control in type 2 diabetes mellitus. Front Immunol, 2023. 14: p. 1125116.

16. Nagakubo, D., et al., DJ-1, a novel oncogene which transforms mouse NIH3T3 cells in cooperation with ras. Biochem Biophys Res Commun, 1997. 231(2): p. 509–13.

17. Liu, F., et al., Mechanisms of DJ-1 neuroprotection in a cellular model of Parkinson’s disease. J Neurochem, 2008. 105(6): p. 2435–53.

18. Zhou, W. and C.R. Freed, DJ-1 up-regulates glutathione synthesis during oxidative stress and inhibits A53T alpha-synuclein toxicity. J Biol Chem, 2005. 280(52): p. 43150–8.

19. Junn, E., et al., Mitochondrial localization of DJ-1 leads to enhanced neuroprotection. J Neurosci Res, 2009. 87(1): p. 123–9.

20. Cuevas, S., et al., Role of nuclear factor erythroid 2-related factor 2 in the oxidative stress-dependent hypertension associated with the depletion of DJ-1. Hypertension, 2015. 65(6): p. 1251–7.

21. Cuevas, S., et al., Role of renal DJ-1 in the pathogenesis of hypertension associated with increased reactive oxygen species production. Hypertension, 2012. 59(2): p. 446–52.

22. Sun, Q., et al., The role of DJ-1/Nrf2 pathway in the pathogenesis of diabetic nephropathy in rats. Ren Fail, 2016. 38(2): p. 294–304.

23. Glat, M.J., et al., Neuroprotective Effect of a DJ-1 Based Peptide in a Toxin Induced Mouse Model of Multiple System Atrophy. PLoS One, 2016. 11(2): p. e0148170.

24. Lev, N., et al., A DJ-1 Based Peptide Attenuates Dopaminergic Degeneration in Mice Models of Parkinson’s Disease via Enhancing Nrf2. PLoS One, 2015. 10(5): p. e0127549.

25. Lev, N., et al., DJ-1 knockout augments disease severity and shortens survival in a mouse model of ALS. PLoS One, 2015. 10(3): p. e0117190.

26. De Miguel, C., et al., ND-13, a DJ-1-Derived Peptide, Attenuates the Renal Expression of Fibrotic and Inflammatory Markers Associated with Unilateral Ureter Obstruction. Int J Mol Sci, 2020. 21(19).

27. Krishnan, S.M., et al., Pharmacological inhibition of the NLRP3 inflammasome reduces blood pressure, renal damage, and dysfunction in salt-sensitive hypertension. Cardiovasc Res, 2019. 115(4): p. 776–787.

28. Wu, M., et al., Inhibition of NLRP3 inflammasome ameliorates podocyte damage by suppressing lipid accumulation in diabetic nephropathy. Metabolism, 2021. 118: p. 154748.

29. Gao, P., et al., NADPH oxidase-induced NALP3 inflammasome activation is driven by thioredoxin-interacting protein which contributes to podocyte injury in hyperglycemia. J Diabetes Res, 2015. 2015: p. 504761.

30. Tshivhase, A.M., T. Matsha, and S. Raghubeer, Resveratrol attenuates high glucose-induced inflammation and improves glucose metabolism in HepG2 cells. Sci Rep, 2024. 14(1): p. 1106.

31. Guo, Q., et al., NF-kappaB in biology and targeted therapy: new insights and translational implications. Signal Transduct Target Ther, 2024. 9(1): p. 53.

32. Iqbal, A., et al., Effect of Hypoglycemia on Inflammatory Responses and the Response to Low-Dose Endotoxemia in Humans. J Clin Endocrinol Metab, 2019. 104(4): p. 1187–1199.

33. Coll, R.C., et al., A small-molecule inhibitor of the NLRP3 inflammasome for the treatment of inflammatory diseases. Nat Med, 2015. 21(3): p. 248–55.

34. Ferreira, N.S., et al., NLRP3 Inflammasome and Mineralocorticoid Receptors Are Associated with Vascular Dysfunction in Type 2 Diabetes Mellitus. Cells, 2019. 8(12).

35. Zhang, C., et al., Activation of Nod-like receptor protein 3 inflammasomes turns on podocyte injury and glomerular sclerosis in hyperhomocysteinemia. Hypertension, 2012. 60(1): p. 154–62.

36. Wada, J. and H. Makino, Innate immunity in diabetes and diabetic nephropathy. Nat Rev Nephrol, 2016. 12(1): p. 13–26.

37. Leu, S.Y., et al., NLRP3 inflammasome activation, metabolic danger signals, and protein binding partners. J Endocrinol, 2023. 257(2).

38. Yong, J., et al., Therapeutic opportunities for pancreatic beta-cell ER stress in diabetes mellitus. Nat Rev Endocrinol, 2021. 17(8): p. 455–467.

39. Shin, J.J., et al., Damage-associated molecular patterns and their pathological relevance in diabetes mellitus. Ageing Res Rev, 2015. 24(Pt A): p. 66–76.

40. Wang, J., et al., High glucose mediates NLRP3 inflammasome activation via upregulation of ELF3 expression. Cell Death Dis, 2020. 11(5): p. 383.

41. Wen, L., et al., ICAM-1 related long noncoding RNA is associated with progression of IgA nephropathy and fibrotic changes in proximal tubular cells. Sci Rep, 2022. 12(1): p. 9645.

42. Kong, H., et al., Targeted P2X7/NLRP3 signaling pathway against inflammation, apoptosis, and pyroptosis of retinal endothelial cells in diabetic retinopathy. Cell Death Dis, 2022. 13(4): p. 336.

43. Ridker, P.M., et al., Inhibition of Interleukin-1beta by Canakinumab and Cardiovascular Outcomes in Patients With Chronic Kidney Disease. J Am Coll Cardiol, 2018. 71(21): p. 2405–2414.

44. Shahzad, K., et al., Nlrp3-inflammasome activation in non-myeloid-derived cells aggravates diabetic nephropathy. Kidney Int, 2015. 87(1): p. 74–84.

45. Hou, Y., et al., NLRP3 inflammasome negatively regulates podocyte autophagy in diabetic nephropathy. Biochem Biophys Res Commun, 2020. 521(3): p. 791–798.

46. Wang, Y., et al., Activation of the NLRC4 inflammasome in renal tubular epithelial cell injury in diabetic nephropathy. Exp Ther Med, 2021. 22(2): p. 814.

47. Song, S., et al., Knockdown of NLRP3 alleviates high glucose or TGFB1-induced EMT in human renal tubular cells. J Mol Endocrinol, 2018. 61(3): p. 101–113.

48. Shahzad, K., et al., Podocyte-specific Nlrp3 inflammasome activation promotes diabetic kidney disease. Kidney Int, 2022. 102(4): p. 766–779.

49. Williams, B.M., et al., The Role of the NLRP3 Inflammasome in Mediating Glomerular and Tubular Injury in Diabetic Nephropathy. Front Physiol, 2022. 13: p. 907504.

50. Caceres, L., et al., Molecular mechanisms underlying NLRP3 inflammasome activation and IL-1beta production in air pollution fine particulate matter (PM(2.5))-primed macrophages. Environ Pollut, 2024. 341: p. 122997.

51. Chow, F.Y., et al., Monocyte chemoattractant protein-1 promotes the development of diabetic renal injury in streptozotocin-treated mice. Kidney Int, 2006. 69(1): p. 73–80.

52. Bohle, A., et al., Human glomerular structure under normal conditions and in isolated glomerular disease. Kidney Int Suppl, 1998. 67: p. S186–8.

53. Uehara-Watanabe, N., et al., Direct evidence of proximal tubular proliferation in early diabetic nephropathy. Sci Rep, 2022. 12(1): p. 778.

54. Hou, Y., et al., CD36 promotes NLRP3 inflammasome activation via the mtROS pathway in renal tubular epithelial cells of diabetic kidneys. Cell Death Dis, 2021. 12(6): p. 523.

55. Solini, A. and I. Novak, Role of the P2X7 receptor in the pathogenesis of type 2 diabetes and its microvascular complications. Curr Opin Pharmacol, 2019. 47: p. 75–81.

56. Glas, R., et al., Purinergic P2X7 receptors regulate secretion of interleukin-1 receptor antagonist and beta cell function and survival. Diabetologia, 2009. 52(8): p. 1579–88.

57. Wang, D., et al., P2X7 receptor mediates NLRP3 inflammasome activation in depression and diabetes. Cell Biosci, 2020. 10: p. 28.

58. Qian, C., et al., P2X7R/AKT/mTOR signaling mediates high glucose-induced decrease in podocyte autophagy. Free Radic Biol Med, 2023. 204: p. 337–346.

59. Lee, S.B. and R. Kalluri, Mechanistic connection between inflammation and fibrosis. Kidney Int Suppl, 2010(119): p. S22–6.

60. Anders, H.J. and M. Ryu, Renal microenvironments and macrophage phenotypes determine progression or resolution of renal inflammation and fibrosis. Kidney Int, 2011. 80(9): p. 915–925.

61. Duffield, J.S., Macrophages and immunologic inflammation of the kidney. Semin Nephrol, 2010. 30(3): p. 234–54.

62. Cao, Q., D.C. Harris, and Y. Wang, Macrophages in kidney injury, inflammation, and fibrosis. Physiology (Bethesda), 2015. 30(3): p. 183–94.

63. Kisseleva, T. and D.A. Brenner, Fibrogenesis of parenchymal organs. Proc Am Thorac Soc, 2008. 5(3): p. 338–42.

64. Brenmoehl, J., et al., Transforming growth factor-beta 1 induces intestinal myofibroblast differentiation and modulates their migration. World J Gastroenterol, 2009. 15(12): p. 1431–42.

65. Wang, L., et al., TGF-Beta as a Master Regulator of Diabetic Nephropathy. Int J Mol Sci, 2021. 22(15).

66. Patibandla, C., et al., Inhibition of glycogen synthase kinase-3 enhances NRF2 protein stability, nuclear localisation and target gene transcription in pancreatic beta cells. Redox Biol, 2024. 71: p. 103117.

67. Li, W., et al., DJ-1 preserves ischemic postconditioning-induced cardioprotection in STZ-induced type 1 diabetic rats: role of PTEN and DJ-1 subcellular translocation. Cell Commun Signal, 2024. 22(1): p. 252.

68. Mulholland, D.J., et al., PTEN and GSK3beta: key regulators of progression to androgen-independent prostate cancer. Oncogene, 2006. 25(3): p. 329–37.

69. Vasconcelos, D.P., et al., 3D chitosan scaffolds impair NLRP3 inflammasome response in macrophages. Acta Biomater, 2019. 91: p. 123–134.

70. Chow, F.Y., et al., Macrophages in streptozotocin-induced diabetic nephropathy: potential role in renal fibrosis. Nephrol Dial Transplant, 2004. 19(12): p. 2987–96.

71. Caballero-Herrero, M.J., et al., Role of Damage-Associated Molecular Patterns (DAMPS) in the Postoperative Period after Colorectal Surgery. Int J Mol Sci, 2023. 24(4).

