## Supplementary material for "ND-13, a DJ-1 derived peptide, as a novel pharmacological approach in the prevention of NLRP3 inflammasome activity in Diabetic Nephropathy": Text of supplementary figures

**Supplementary data**

**Supplementary figure 1.** **IL-1β release by bone marrow derived macrophages in STZ-induced diabetic C57BL/6 mice and control C57BL/6 mice**. **(a)** Release of IL-1β in control mice after none or different stimuli. **(b)** Release of IL-1β in STZ-induced diabetic mice after none or different stimuli. **(c)** Release of IL-1β in STZ-induced diabetic mice treated with ND-13 after none or different stimuli. **(d)** Release of IL-1β in STZ-induced diabetic mice treated with MCC950 after none or different stimuli. Data is represented as mean ± SEM; t-test two-sided was used to compare with resting column in A-D. Significance levels are indicated as follow *****p<0.0001*, ***p<0.01*, **p<0.05*. Control n=8, STZ n=10, STZ+ND-13 n=8, STZ+MCC950 n=8.

**Supplementary figure 2. (a)** Protein expression of Synaptopodin in whole kidney (representative n=3 from each group of mice). **(b)** Protein expression of Desmin in whole kidney (representative n=3 from each group of mice) **(c)** Protein expression of GSK-3 in bone marrow derived macrophages (n=3). Data is represented as mean ± SEM; Tukey's multiple comparisons test was used in a-c.

**Supplementary figure 3. High glucose concentration does not affect NLRP3 activation in vitro**. **(a)** Release of Il-1β in the culture medium. **(b)** Release of TNF-α in the culture medium. **(c)** Percentage of cell death compared to positive control. Data is represented as mean ± SEM (n=3); t-test two-sided was used in a-c to compare between the group with high glucose (white column) and its corresponding group with low glucose (grey column). Significance levels are indicated as follow: ***p<0.005*, **p<0.05*.
